## Supplementary Material for "Whole-cell modeling in yeast predicts compartment-specific proteome constraints that drive metabolic strategies"

**This document includes:**

**Supplementary Figures 1-12**

**Supplementary Notes**

**Supplementary Tables and Datasets (separate .xlsx files)**

**Table S1.** Protein-pathway and protein-compartment mappings, used in the study.

**Table S2.** Lists of genes, activated by PKA-, TOR-, or cooperative PKA and TOR signalling.

**Dataset S1.** Flux and yield data of yeast cultures, grown in glucose-limited chemostats, excess sugar batches and in glucose-excess batches treated with cycloheximide.

**Dataset S2.** Label-free quantitative proteomics data for the cultures, grown in glucose-limited chemostats.

**Dataset S3.** Label-free quantitative proteomics data for the cultures, grown in excess sugar batches.

**Dataset S4.** Label-free quantitative proteomics data for the cultures, grown in excess glucose batches, treated with cycloheximide.

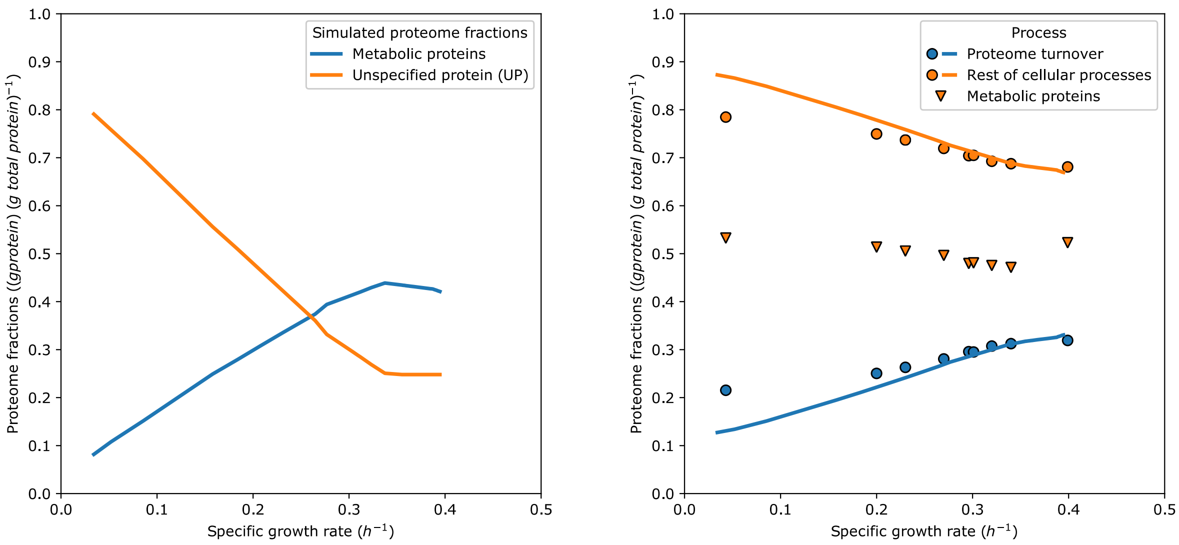

**Fig. S1.** Computed protein fractions as a function of growth rate. Left, predicted fractions of the unspecified protein (orange) and metabolic proteins (blue); right, fractions of proteome turnover-associated proteins (blue) and proteins, participating in other cellular processes (orange).

The dots represent experimentally determined proteome fractions from glucose-limited chemostats (at $D = 0.20$ to $0.34\, h^{-1}$) and trehalose- and glucose-excess batch cultivations ($\mu = 0.043$ and $\mu = 0.399\, h^{-1}$; left-most and right-most points, respectively), and lines are model predictions in glucose-limited conditions. The annotations for the "Proteome turnover" class were collected by the Authors (Table S1 of the Manuscript); for "Metabolic proteins", we used KEGG ontology annotations for processes related to metabolism.

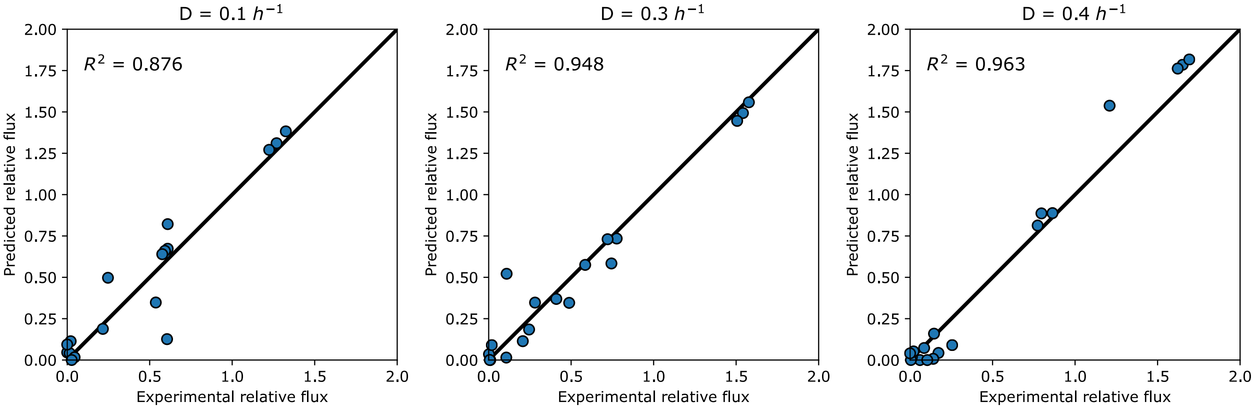

**Fig. S2.** Comparison of experimentally determined (abscissa) and predicted (ordinate) fluxes in central carbon metabolism of glucose-limited *S. cerevisiae*. Simulations of glucose-limited growth were performed by varying the hexose transporter saturation, as for the Figure 2d of the main text. Experimental data for the intracellular fluxes at three dilution rates ($D = 0.1$, $0.3$ and $0.4\ h^{-1}$) were taken from (Frick & Wittmann, 2005). Flux values were normalized to those of the hexokinase reaction.

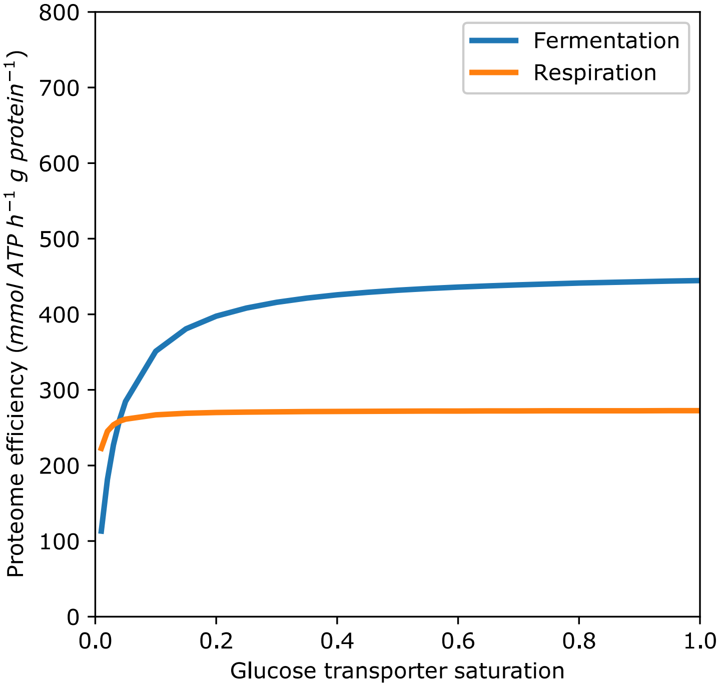

**Fig. S3.** Dependence of the proteome efficiency (measured in $mmol\ ATP/g\ protein/h$) for energy harvesting from glucose through respiration and fermentation pathways on the saturation of the glucose transporter. Computations were performed according to (Chen & Nielsen, 2019). In brief, we compiled information on the molecular weights of the proteins that participate in glycolysis, TCA cycle and oxidative phosphorylation, together with respective turnover values. Then we computed the amounts of each protein (in grams) needed in order to sustain a flux of $1\,mmol\ ATP/h$ using either respiratory of fermentative pathway under different saturation of glucose transporter. The final value of the proteome efficiency was computed by evaluating the flux of ATP production per gram of protein invested.

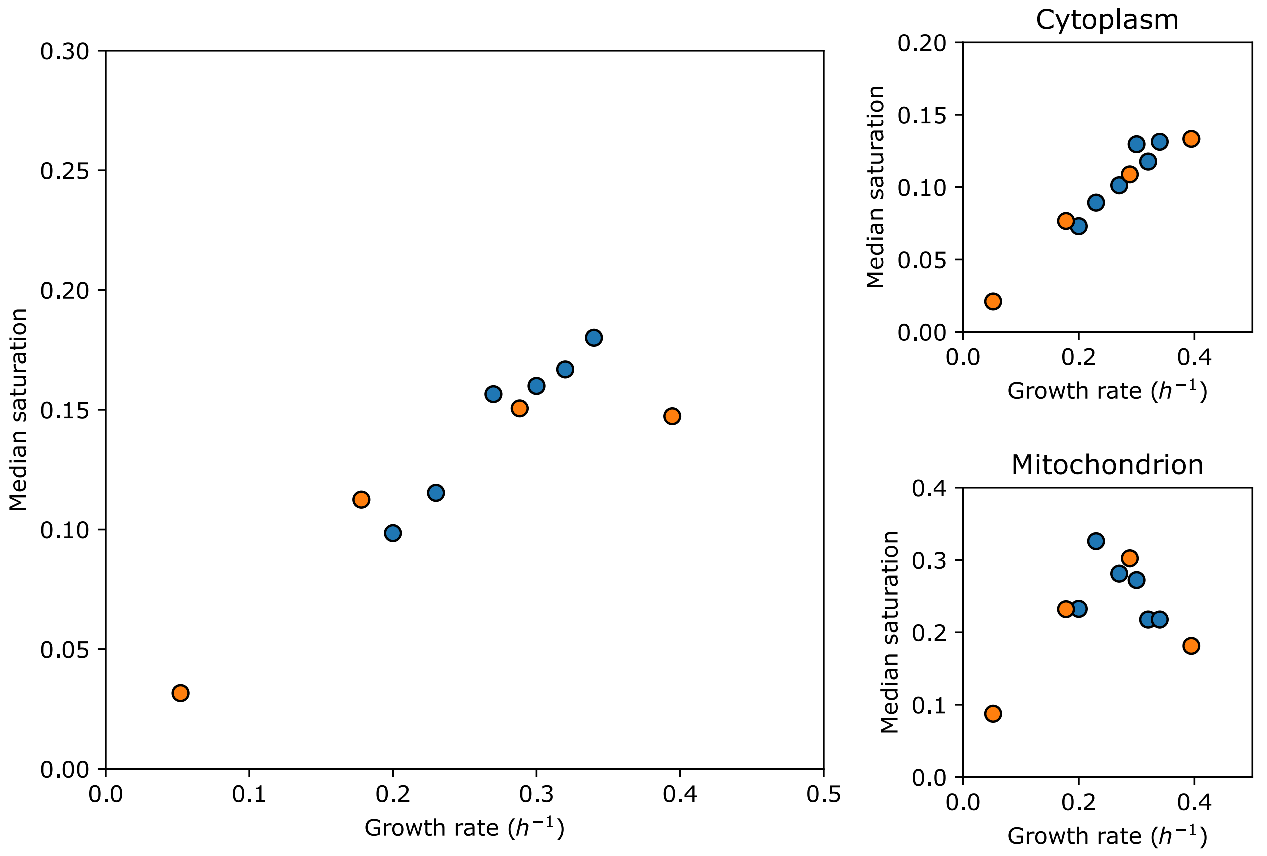

**Fig. S4.** Growth-rate dependence of the median saturation factor of metabolic enzymes. Blue points, glucose-limited chemostat cultures; orange points, sugar excess cultures. The saturation factor was determined as a ratio between the expression level predicted by the pcYeast model vs. the experimentally determined expression level from quantitative proteomics measurements. Ratios were taken into consideration only if the protein abundance was both present and nonzero in both datasets. All proteins with a saturation factor higher than 1 were assigned a factor of 1.1.

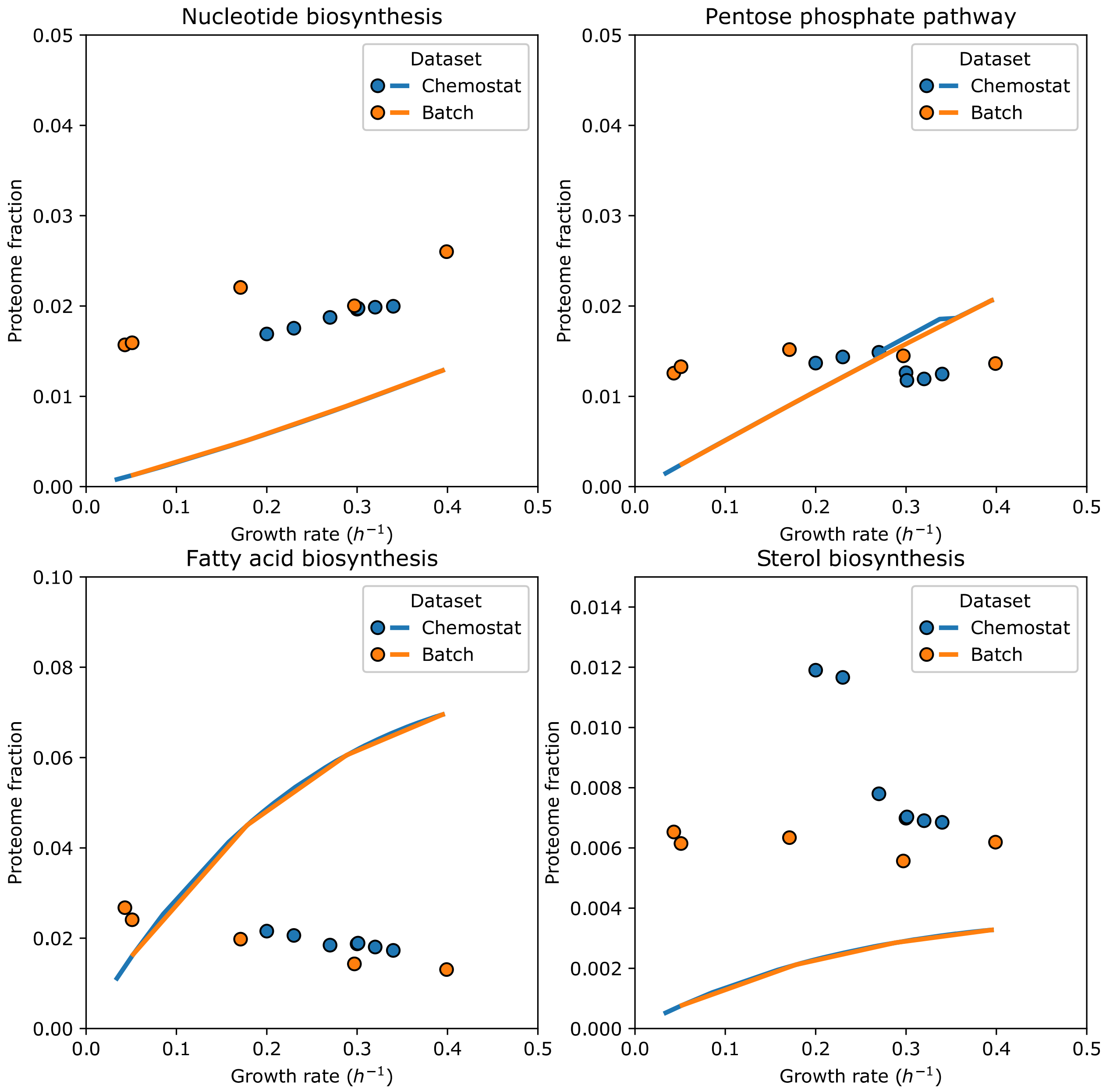

**Fig. S5.** Comparison of experimentally determined (circles) and predicted (lines) proteome fractions occupied by proteins in some biosynthetic pathways). Pathway-protein mappings (Table S1) were manually curated, using the reference proteome of *S. cerevisiae* from the UniProt database.

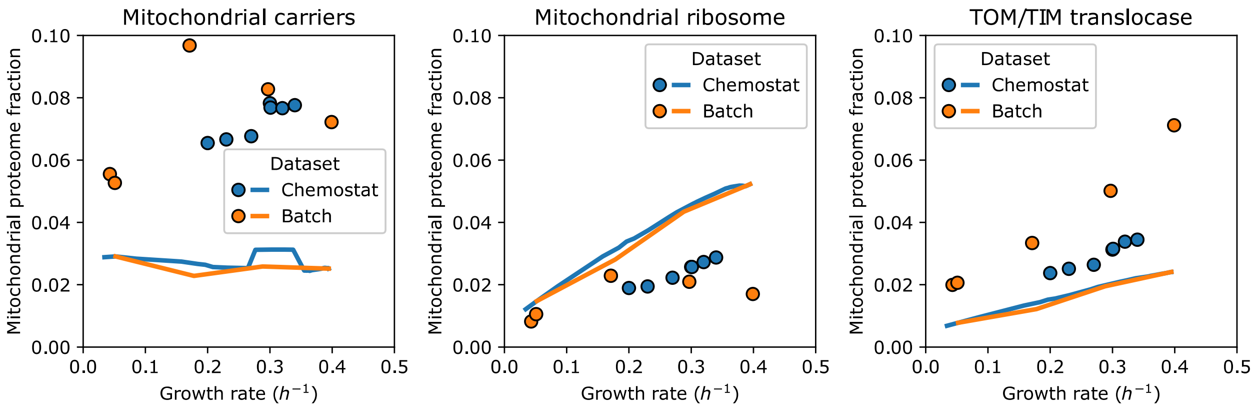

**Fig. S6.** Comparison of experimentally determined (circles) and predicted (lines) *mitochondrial* proteome fractions (*i.e.* protein per total mitochondrial protein) occupied by proteins in pathways, located in mitochondria and mitochondrial membrane. Pathway-protein mappings (Table S1) were manually curated, using the reference proteome of *S. cerevisiae* from the UniProt database.

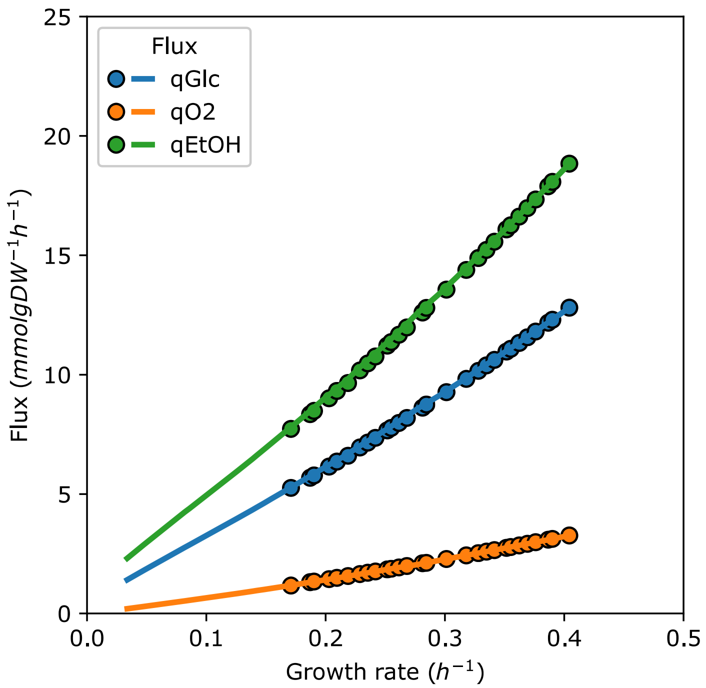

**Fig. S7.** Predicted metabolic fluxes for dose-dependent mCherry overexpression in glucose minimal medium (simulation performed analogously to the simulation of mCherry overexpression in SD medium, Fig. 1D, points) and inhibition of translation (lines). The overexpression of mCherry was simulated by setting the desired proteome mass fraction, which mCherry should occupy and running the binary search algorithm to determine the highest growth rate which would still result in a feasible linear program. For inhibition of ribosomes, same procedure was applied, but instead of pre-setting the level of gratuitous protein expression, the activity factor of ribosomes ($0 < f_{Ribo}^{A} \leq 1$) was varied (see Ribosome Capacity Constraint 1).

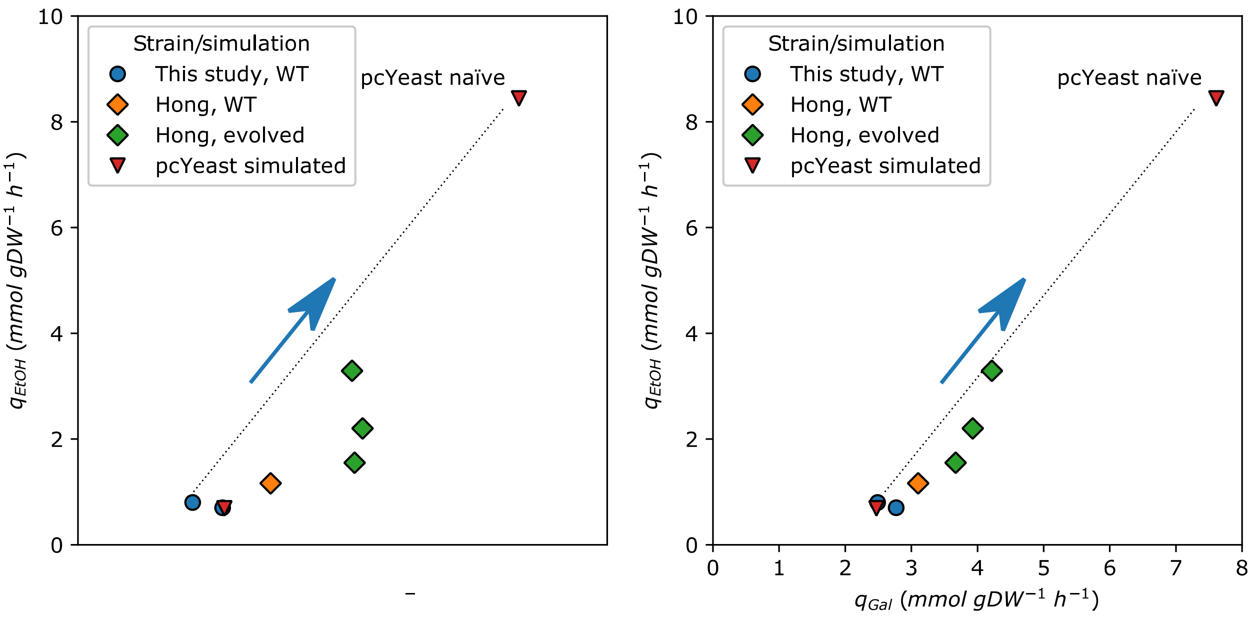

**Fig. S8.** Comparison of the physiology of wild-type and galactose-evolved strains of S. cerevisiae and pcYeast simulations. Left, relationship between specific growth rate and ethanol formation; right, relationship between consumption of galactose and ethanol formation. In both our study and (Hong et al., 2011), the WT strain was CEN.PK113-7D. The "naïve" simulation of pcYeast corresponds to the maximal achievable growth rate on galactose without parameter tweaking (see Supplementary Notes for more details).

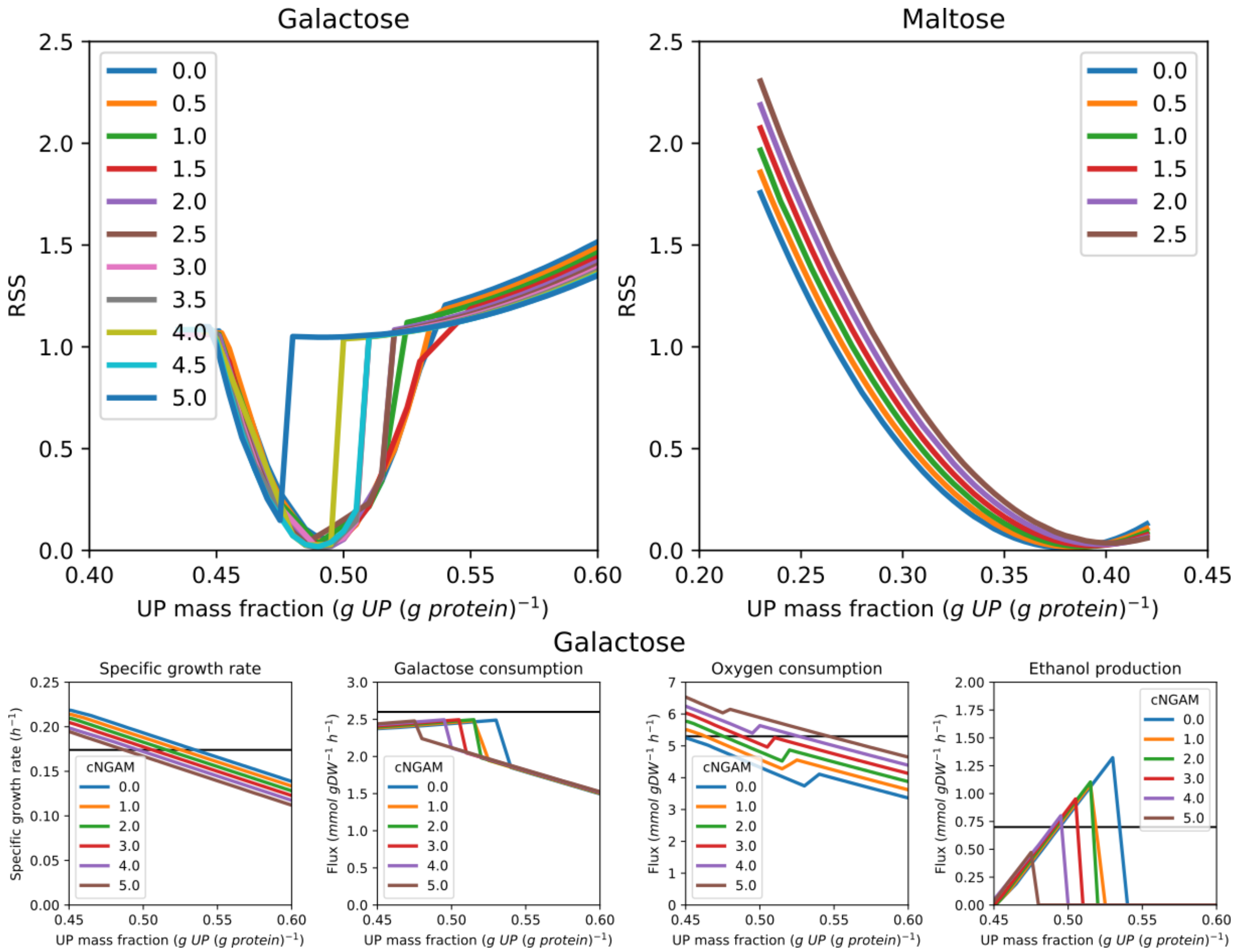

**Fig. S9.** Estimation of the parameter values which best represent the experimentally measured physiology of galactose- and maltose-excess batch cultures. The parameters scanned are the minimal UP fraction together with the carbon source-associated NGAM (cNGAM) (upper panels). Prediction of the specific growth rate and metabolic fluxes for different combinations of parameters for galactose are provided in the bottom panels. Details of the procedure used to estimate the parameters are provided in Supplementary Notes.

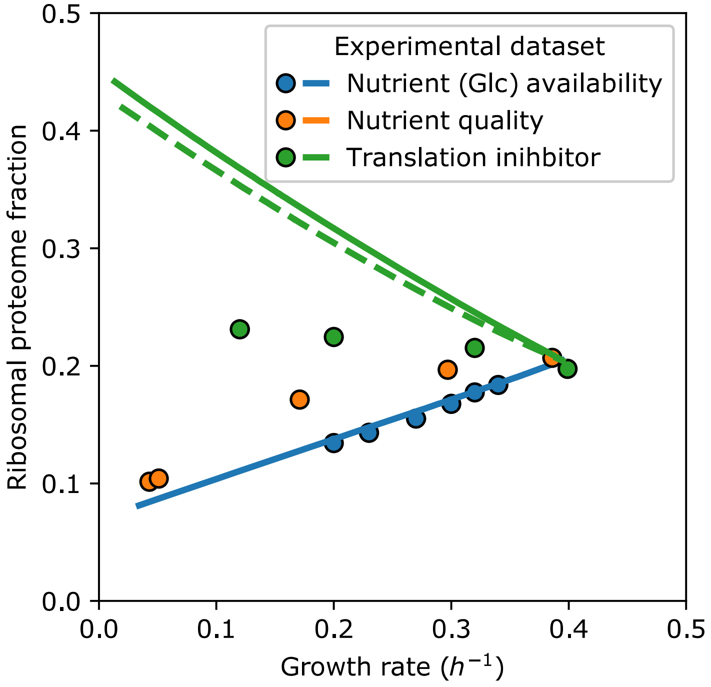

**Fig. S10.** Ribosomal proteome fractions (g r-protein/g protein) in different growth scenarios. The dashed line for the simulation of translation inhibition with cycloheximide represents the case where the fraction of inactive ribosomes $\phi_R^0$ (see Main Text for details) was linked to the factor of ribosome inhibition: $\phi_R^0 = \phi_R^0(non - inhibited) \times f_{Ribo}^A$, where $f_{Ribo}^A = 1 - f_{Ribo}^{Inh}$. Lines are model predictions, symbols are experimental data points.

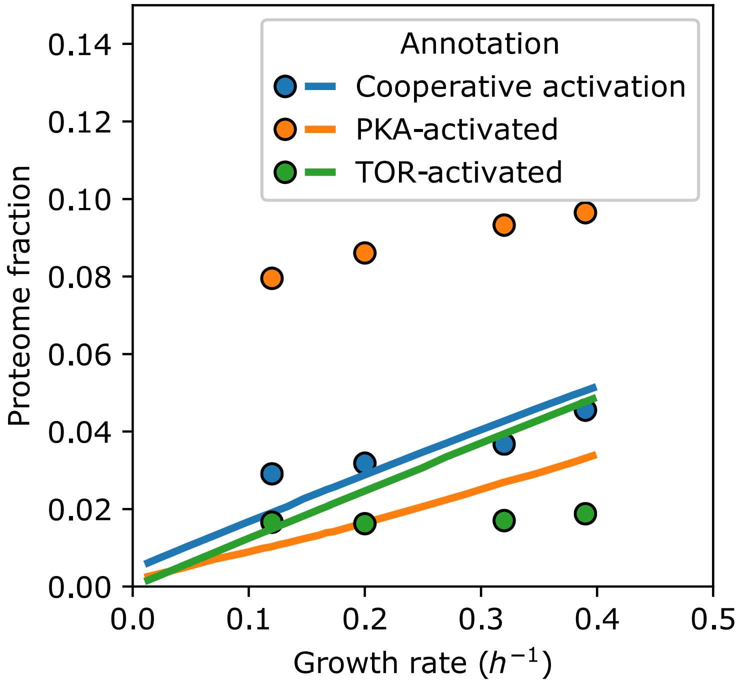

**FigS11.** Experimentally determined (circles) and predicted (lines) proteome fractions of proteins that were activated either by TOR, by PKA, or cooperatively activated by both TOR and PKA signalling, as determined by (Kunkel et al., 2019). Groups of specifically activated genes were identified by considering the expression coefficients computed for *S. cerevisiae*, transferred from glycerol- to glucose-rich media containing inhibitors of either PKA, TOR, or both, 20 minutes post-transfer. We selected the genes, for which the expression component was $\geq 2$ (Table S2).

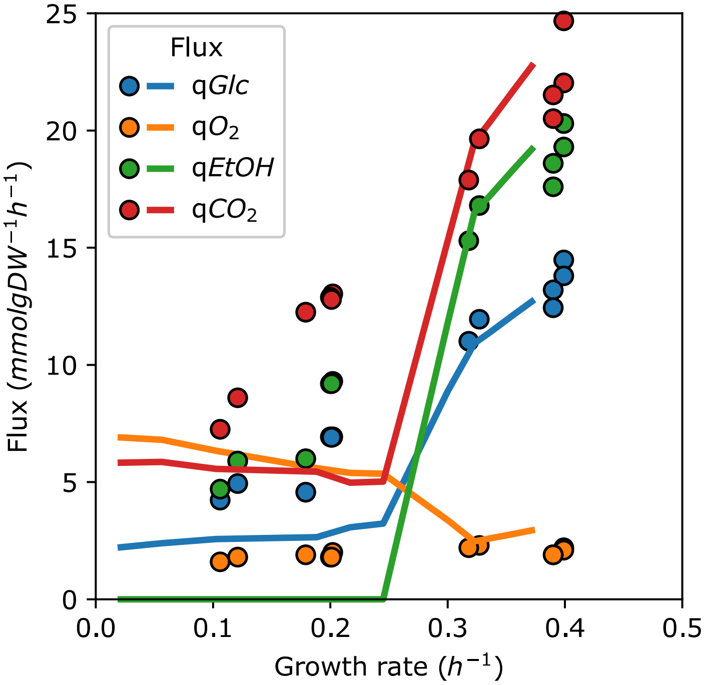

**Fig. S12.** The effect of translation inhibition by cycloheximide on the main catabolic fluxes in controlled aerobic batch fermentations on glucose with a phenomenological constraint on the ribosomal fraction. Lines are model predictions with a forced constraint on the maximal ribosomal proteome fraction of $0.25\ g\ r\ protein/\ g\ protein$, symbols are experimental data points.

**Supplementary Notes**

**Generation and Formulation of the pcYeast**

### 1. Formulation of the Linear Program (LP)

min $v_{Dummy}$, subject to:

1. **Mass balance constraints**:
   $S \times \boldsymbol{v} = 0,$
   where $S$ is the stoichiometry matrix connecting all metabolites to reactions, and $\boldsymbol{v}$ is the flux vector for all reactions.
2. **Fixed growth rate at every iteration**: $\mu = \mu_{set}$.
3. **Enzyme capacity constraints**:
   $v_i = k_{cat,i}\, [e_i]\, f_i,\ i \in metabolic\ enzymes$
   where $[e_i]$ is the concentration of a metabolic enzyme $e_i$, $k_{cat,i}$ is the catalytic rate of enzyme $e_i$, and $f_i$ is the saturation function of the enzyme $e_i$. Normally, the saturation level depends on enzyme kinetics and metabolite concentration information. We fix the saturation factor $f_i = 1$ for all enzymes except for hexose transporters and thus compute the minimal proteome requirement to sustain the metabolic fluxes.
4. **Protein synthesis, folding and degradation constraints**:
   $\sum_{i \in cytosolic\ ribosome} v_{syn,i} = k_{cat,cytosolic\ ribosome}\ [Ribosomes]$ ,
   $\sum_{i \in mitochondrial\ ribosome} v_{syn,i} = k_{cat,mitochondrial\ ribosome}\ [Mito\ ribosomes]$,
   $\sum_{i \in folded\ by\ j} v_{folding,i} = k_{folding,j}\ [j]\ ,\ j \in chaperones,$
   $\sum_{i \in degraded\ by\ j} v_{degradation,i} = k_{degradation,j}\ [j]\ ,\ j \in proteases,$
   These constraints link the protein life-cycle processes to the availability of appropriate components.
5. **Compartment-specific proteome constraints**:
   $\sum v_{syn,i}\, w_i \le V_{compartment}$ or $\sum v_{syn,i}\, w_i \le A_{compartment}$, $i \in compartment,$
   which limit the proteome according to the volume or area of the appropriate compartment.
6. **Total proteome constraint**:
   $\sum v_{syn,i}\, w_i = P(\mu),\ i \in proteins$
   defining growth rate-dependent protein content of the dry weight of cells (in $g\ protein/gDW$). Note that this is an equality constraint, and has to be fulfilled also at low growth rates, where metabolic fluxes and corresponding protein demands are low. We use an unspecified protein $UP$ with average protein composition to maintain this equality and thus protein density.
7. **Growth-related metabolic reactions**:
   $v_i = k \times \mu$
   Examples of such reactions are the growth-associated ATP maintenance (GAM), which accounts for the ATP expenditures coupled to growth that are not explicitly modeled. The biomass reaction describes the demand for all biomass components (except proteins, which are explicitly modeled as described above), proportional to the growth rate.

### 2. Model Constraints and Implementation

We converted the linear program of our model to CPLEX LP format (.lp) and used the SoPlex solver (Gleixner et al., 2017) to solve the problem with high precision (maximal feasibility and optimality violation: $10^{-12}$). The advantage of writing our model as CPLEX LP format is the ability to add constraints directly without any restrictions imposed by the software which is used to handle genome-scale models. Additionally, most solvers read CPLEX LP format, so we are not restricted to the solvers offered by RAVEN (Wang et al., 2018), COBRA (Heirendt et al., 2019) or PySCeS CBMPy (Olivier et al., 2005) packages. The maximal feasible growth rate $\mu_{max}$ is determined by using binary search, i.e. running LPs to test for feasibility in a predefined range of growth rates to find the highest feasible value (see Section 1). We consider the simulated growth rate to be the highest feasible ($\mu_{max}$) if a very small increment in growth rate $\Delta\mu$ results in an infeasible linear program (i.e. at $\mu = \mu_{max} + \Delta\mu$) . We set $\Delta\mu \leq 10^{-4}\ h^{-1}$ in our simulations. Note that the model implementation is available in two formats, based in either MATLAB or Python, and the underlying logic/assumptions are valid for both implementations.

The model introduces novel proteome constraints which are in essence phenomenological limits based on observation on the CEN.PK113-7D strain of *S. cerevisiae*, rather than being hard physico-chemical constraints. In the next subsections, we describe every constraint in the model in more detail.

#### 2.1 Mass balance constraints

**Mass Balance Constraint 1:**

The first constraint in our model is the steady-state condition ($S \times \boldsymbol{v} = 0$), where $\boldsymbol{v}$ is the flux vector and $S$ is the stoichiometric matrix of $i \times j$ columns (reactions) and rows (metabolites), respectively. For any reversible reaction in the Yeast7.6 model, we split them into two unidirectional reactions, forward and reverse. This constraint is the main constraint in stoichiometric models, and is applies to all molecular species in the cell. In the constraint expressions below, we use the convention of $v_i$ for a flux.

#### 2.2 Enzyme capacity constraints

**Flux Capacity Constraint 1: Metabolic coupling**

For all metabolic reactions $\{v_1, v_2, \dots, v_n\} \in metabolic\ rxns$ that are catalyzed by enzymes (with enzyme concentrations $[e_i]$), the model now has a demand of producing metabolic enzymes. Moreover, every enzyme $e_i$ is diluted by growth and/or degraded towards individual amino acids. In order not to violate the steady state of the system, the sum of dilution and degradation fluxes must be equal to that of protein synthesis. Thus we define the steady-state enzyme concentration as follows:

$$[e_i] = \frac{v_{syn,i}}{\mu + k_{deg}}$$

where $k_{deg}$ is the protein degradation rate. Assuming the enzyme works at its maximal reaction rate, i.e. $v = v_{max}$, we determine the relationship between metabolic fluxes and enzyme demand (Pavlov & Ehrenberg, 2013):

$$[e_i] = \frac{v_{max}}{k_{cat,i}}$$

where $k_{cat,i}$ is the turnover constant for the enzyme $e_i$.

We fixed the $k_{deg}$ for all peptides in the model to the median protein degradation rate $k_{deg} = 0.043\ h^{-1}$, according to experimental measurements of protein stability at $\mu = 0.1\ h^{-1}$ (Lahtvee et al., 2017). To establish the ratio between fluxes through metabolic reactions and enzyme demands, we acquired $k_{cat,i}$ values using the GECKO toolbox (Sánchez et al., 2017), as well as manually curated values for enzymes in central carbon metabolism (Nilsson & Nielsen, 2016), or other sources, such as BRENDA (Placzek et al., 2017) or SABIO-RK (Wittig et al., 2018).

In the model, the costs for both monomeric enzymes, as well as their complexes, are associated with complex assembly reactions. Enzyme complexes are formed based on the known stoichiometric ratios of the individual subunits:

$$R_{complex\ formation,i}\colon \sum_{k \in cplx_i} s_k e_k \rightarrow cplx_i$$

where $s_k$ is the stoichiometric coefficient of the subunit $e_k$. We collected information about stoichiometry of the complexes from UniProt (Bateman et al., 2017), MetaCyc (Caspi et al., 2020), and Protein Data Bank (Burley et al., 2018). For each complex of metabolic enzymes, its demand is considered to be the sum of costs for individual reactions, using this enzymatic complex:

$$\sum_{j=1}^{m} \frac{v_j}{k_{cat,i,j}} \leq \frac{v_{complex\ formation,i}}{\mu + k_{deg}}$$

where $k_{cat,i,j}$ is the turnover constant of the complex $i$, catalyzing the $j$-th reaction.

#### Protein Balance Constraint 2a: Dilution by growth

All proteins $e_i$ are diluted by growth, i.e. with a rate constant equal to the growth rate $\mu$. Therefore, we set the flux through the dilution reaction of the complex $cplx$ to $\mu \times [e_i]$:

$$R_{dil,i}\colon cplx_i \rightarrow \emptyset \text{ and}$$

$$v_{dil,i} - \mu \frac{v_{complex\ formation,i}}{\mu + k_{deg}} = 0.$$

#### Protein Balance Constraint 2b: Degradation to free amino acids

Similarly, we set the flux through degradation reaction of the complex $cplx$ to $k_{deg} \times [e_i]$.

$$R_{deg,i}\colon cplx_i + mn\ ATP + (m + 2n - 1)\ H_2O \rightarrow \sum_{j \in aa} kj + mn\ ADP + mn\ P_i + mn\ H^+, \sum j = n \text{ and}$$

$$v_{deg,i} - k_{deg} \frac{v_{complex\ formation,i}}{\mu + k_{deg}} = 0.$$

#### 2.3 Ribosome, chaperone and protease constraints

**Ribosome Capacity Constraint 1: Capacity of the cytosolic ribosome**

We considered the ribosome demand to be affected by the processes of ribosome scanning, translation initiation, elongation and termination. We implemented expenditures of resources for the processes outside translation elongation as respective increments to the length of the amino acid chain. We constructed a coupling constraint for the synthesis reaction of peptide species $e_i$ as follows: the demand of ribosomes for synthesis of $e_i$ (denoted as $Ribo_{e_i}$) is dependent on peptide elongation rate $k_{cat,ribo}$, as well as on the length of the peptide $l_{peptide}$ (i.e. translation elongation), together with the increments of $l_i$ and $l_{scanning}$ for translation initiation, number of scanning steps through the 5'UTR region of the transcript for the protein $e_i$ and translation termination (an equivalent of one amino acid), respectively. For each peptide species $e_i$, the constraint is defined as:

$$v_{syn,e_i}\,[mmol/gDW/h] = \frac{k_{cat,ribo}\,[aa/h]}{l_{i,e_i} + l_{scanning,e_i} + l_{peptide,e_i}\,[aa] + 1} Ribo_{e_i}\,[mmol/gDW]$$

where $k_{cat,ribo}$ is the maximal peptide elongation rate of $k_{cat,ribo} = 10.5\,aa\,s^{-1} = 3.78 \times 10^4\,aa\,h^{-1}$ (Metzl-Raz et al., 2017).

Section 6 describes in more detail how we also adjust ribosomal protein synthesis according to estimation that ca. 8% of yeast proteome is occupied by ribosomal proteins at zero growth (Metzl-Raz et al., 2017). Here we provide only the description of how we accounted for this in the model. We defined the extra demand for the ribosomal protein allocation by the following expression:

$$Ribo_{nontranslating} = \phi_R^0 \times f_{P,biomass}(\mu)[g\ protein/gDW]/M_{P,ribo}[g\ protein/mmol]$$

where $\phi_R^0 = 0.053$ is the proteome fraction, always filled with non-translating ribosomes (see Section 6) and $M_{P,ribo} = 1388.488\ g/mmol$ is the molar mass of the protein component of the fully assembled ribosome. Following (Pavlov & Ehrenberg, 2013), we define that $Ribo_{e_i}$ is the amount of ribosomes that translates $e_i$ and thus the total ribosome abundance in the model equals $\sum_{e_i} Ribo_{e_i} + Ribo_{nontranslating} = [Ribosome]$.

We modulated the catalytic activity of all protein translation machinery (i.e. ribosomes, ribosome assembly factors, as well as translation initiation, elongation and termination factors), by the ribosome activity factor, which is $f_{Ribo}^A = 1$ for unperturbed cells. In case of inhibition of ribosomal activity (e.g. cycloheximide treatment), the $1 - f_{Ribo}^{Inh} = f_{Ribo}^A < 1$.

The balance between fluxes through ribosome biosynthesis, dilution, and degradation reactions is derived in the same manner as for the metabolic reactions (see Constraints 1, 2a and 2b in Section 2.2). By using the relationship $\sum_{e_i} Ribo_{e_i} + Ribo_{non-translating} = [Ribosome]$ for summing up the individual amounts of ribosomes, synthesizing every single peptide species to the total ribosomal pool, we set the requirement for ribosome synthesis (assembly of ribosomal units) as follows:

$$[Ribosome] \le f_{Ribo}^A \times \frac{v_{complex\,formation,ribo}}{\mu + k_{deg,ribo}}$$

where $v_{complex\,formation,ribo}$ is the ribosome complex formation (assembly) reaction, for which we used annotations for the subunits of cytosolic ribosome from (de la Cruz et al., 2015).

#### Ribosome Capacity Constraint 2: Cytosolic translation initiation factors

Translation initiation in yeast has been studied in great detail to reveal the mechanism of translation initiation in eukaryotic cells (Hinnebusch, 2014). A number of translation initiation factors ($eIF1A$, $eIF1B$, $eIF2A$, $eIF1B$, $eIF3$, $eIF4A$, $eIF4B$, $eIF4E$, $eIF4G$, $eIF5$) participate in this process.

Some of the initiation factors also form separate complexes which scan the 5'UTR region until finding the first codon AUG to form the 48S ribosomal complex. Thus the 5'UTR length greatly influences the cost of the scanning process in translation initiation. To determine the TSS (translation starting site i.e. number of nucleotides before the coding sequence starts) for each gene, we used SMORE-seq data (Parky et al., 2014). We compared the SMORE-seq data with the yeast genome sequence R64-1-1 from the SGD database (Engel et al., 2014) to determine the start and end position of the CDS. For a gene with no experimental estimation of TSS, we assume its 5'UTR length to be 50 nucleotides (median of 5'UTR length). For a gene that has TSS inside the CDS, we assume its 5'UTR length to be zero. This information is then used to compute the mRNA-specific costs of translation initiation.

To describe the demand for translation initiation factors, we defined all initiation translation factors as a single complex ($TIF$). One of the increments to the expenses to the protein synthesis, as described before, is the time for the ribosome to scan along the mRNA to find the first coding triplet. (Berthelot et al., 2004) proposed a random walk algorithm to estimate the number of steps in the 5'UTR region ($l_{scanning}$) (see Ribosome Capacity Constraint 1). The complex hydrolyzes one ATP per each scanning step.

Here, we assumed that a ribosome scans transcripts at the rate of about 10 nucleotides per second ($k_{scan} = 10\ nt\ s^{-1} = 3.6 \times 10^3\ nt\ h^{-1}$), which is close to the maximal peptide elongation rate of $k_{cat,ribo} = 10.5\ aa\ s^{-1}$. Thus, from the perspective of ribosomal demand, we considered one scanning step at the 5'UTR region as one additional amino acid and added the number of scanning steps ($l_{scanning}$) to the total protein length. For each peptide species $e_i$ we define the relationship between translation of protein species $e_i$ and the demand of $TIF$ as follows:

$$v_{syn,e_i}\,[mmol/gDW/h] - \frac{k_{scan}[nt\ h^{-1}]}{l_{scanning,i}\ [nt]} TIF_{e_i}[mmol/gDW/h] = 0$$

where $v_{syn,e_i}$ is a flux through the synthesis reaction of $e_i$, $k_{scan}$ is the ribosome scanning rate and $TIF_{e_i}$ is the amount of $TIF$ required for initiation of synthesis of peptide $e_i$.

Alike the ribosomal amounts for each peptide species $e_i$, the amounts of $TIF$ could be summed up to determine the total pool of $TIF$: $\sum_{e_i} TIF_{e_i} = [Translation\ initiation\ factors]$. We thus set the requirement for the synthesis of $TIF$ (assembly of translation initiation factors into the $TIF$ complex) as follows:

$$[Translation\ initiation\ factors] \le f_{Ribo}^{A} \times \frac{v_{complex\,formation,TIF}}{\mu + k_{deg,TIF}}.$$

#### Ribosome Capacity Constraint 3: Cytosolic translation elongation factors

We also accounted for the demand of translation factors $eEF1A$, , $eEF1B$, $eEF2$, $eEF3$ (Andersen et al., 2006; Chakraburtty, 2001; Dever & Green, 2012), as well as the hydrolysis of the nucleotide triphosphates during peptide translation. In total, 2 GTP and 2 ATP are spent for elongation of the peptide chain by one amino acid.

The elongation factors are required to catalyze their reactions at the speed equivalent to the peptide elongation, thus we assumed $k_{cat,EF} = k_{cat,ribo}$. To couple the elongation translation factor (i.e., $\{eEF1A, eEF1B, eEF2, eEF3\}$) biosynthesis reactions with the demand of these factors in translation of peptides, we considered that synthesis of each protein $e_i$ requires each of the $eEF_j$ ($\mathrm{eEF}_{j,e_i}$). For each peptide species $e_i$ and $eEF_j$, the constraint was defined as follows:

$$v_{syn,e_i} - \frac{k_{cat,EF}}{l_{peptide,e_i}} \, eEF_{j,e_i} = 0$$

where $v_{syn,e_i}$ is a flux through protein synthesis reaction of $e_i$ and $l_{peptide,e_i}$ is the length of protein $e_i$. Alike the ribosomal amounts for each peptide species $e_i$, the amounts of $eEF_{j,e_i}$ could be summed up to determine the total pool of $eEF_j$:

$$\sum_{e_i} eEF_{j,e_i} = [eEF_j]$$

By using this relationship, we set the requirement for the complex formation of $eEF_j$ as follows:

$$[eEF_j] \leq f_{Ribo}^{A} \times \frac{v_{complex\ formation,eEF_j}}{\mu + k_{deg,eEF_j}}.$$

#### Ribosome Capacity Constraint 4: Cytosolic translation termination factors

We adopted a termination model proposed by (Dever & Green, 2012). In this model, the translation termination factor ($eRF1$) binds with GTP-bound form of $eRF3$. $eRF3 - GTP$ is a ribosome-binding factor that recognizes the stop codon. Thus, translation termination requires one molecule of GTP.

To also account for the cost of translation termination factors, we set the demand of eRF ($eRF_{\ e_i}$) for each protein synthesis reaction. We considered the termination rate to be equal to $k_{cat,eRF} = 0.15\ s^{-1}$ (Shoemaker & Green, 2011). Thus we formulated the constraint as follows:

$$v_{syn,e_i} - k_{cat,\mathrm{eRF}} \, \mathrm{eRF}_{\ e_i} = 0$$

where $v_{syn,e_i}$ is a flux through protein synthesis reaction of $e_i$ and $k_{cat,\mathrm{eRF},}$ is the turnover rate of $eRF2$ binding. Alike the ribosomal amounts for each peptide species $e_i$, the amounts of $eRF$ could be summed up to determine the total pool of $eRF$:

$$\sum_{e_i} eRF_{\ e_i} = [eRF]$$

By using this relationship, we set the requirement for the complex formation of $eRF$ as follows:

$$[eRF] \leq f_{Ribo}^{A} \times \frac{v_{complex\ formation,eRF}}{\mu + k_{deg,eRF}}.$$

**Ribosome Capacity Constraint 5: Cytosolic ribosome assembly factors**

Ribosome assembly factors are required for assembling new ribosome from its subunits. Assembly factors facilitate assembly of ribosomal subunits into supercomplexes, which are transported from nucleus to cytoplasm. After that, the ribosomal subunits go through maturation. We used our proteomics data of sugar-excess cultures in order to estimate the turnover number of the ribosome assembly factors, so the predicted demand of these factors would match the experimentally determined abundance. Following that, we set the $k_{cat,ribo\ assembly} = 1\ min^{-1}$.

As there are many species of ribosome assembly factors, we consider them as a single complex and assume that all ribosomal assembly factors are bound to ribosomal subunits (Woolford, John Jr, Baserga, 2013). We thus introduced a coupling constraint, setting the demand for ribosome assembly factors $v_{complex\ formation,ribo\ assembly}$ to be proportional to the ribosome assembly reaction $v_{complex\ formation,ribo}$ as follows:

$$v_{complex\ formation,ribo} \leq f_{Ribo}^{A} \times k_{cat,ribo\ assembly} \frac{v_{complex\ formation,ribo\ assembly}}{\mu + k_{deg,ribo\ assembly}}.$$

**Ribosome Capacity Constraint 6: Mitochondrial ribosome capacity**

The mitochondrial ribosome translates the peptides for 7 proteins of oxidative phosphorylation system and one subunit of the mitochondrial ribosome. Due to this fact, we provided a simplified description of the mitochondrial ribosome in the model, i.e. we did not take into account the expenses of translation initiation, termination and/or associated factors. Even though, we modelled the translation elongation in a similar fashion as for the cytosolic ribosome (Ribosome Capacity Constraint 1). With defining the demand for the mitochondrial ribosome $MRibo_{e_i}$ for the synthesis of the peptide $e_i$ with amino acid chain length $l_{peptide,e_i}$, we formulated the following constraint:

$$v_{syn,mito,e_i} = \frac{k_{cat,ribo,mito}}{l_{peptide,e_i}} MRibo_{e_i}$$

where the elongation rate of the mitochondrial ribosome was estimated to be $k_{cat,ribo,mito} = 2.7\ aa\ s^{-1}$, according to the proteomics measurements in glucose-limited chemostat cultures.

By using the relationship $\sum_{e_i} MRibo_{e_i} = [mitoRibosome]$ between the amount of mitochondrial ribosomes, synthesizing single peptide species and total pool of mitochondrial ribosomes, we set the requirement for ribosome synthesis (assembly of ribosomal units) as follows:

$$[mitoRibosome] \leq \frac{v_{complex\ formation,mitoribo}}{\mu + k_{deg,mitoribo}}$$

where $v_{complex\ formation,mitoribo}$ is the mitochondrial ribosome complex formation (assembly) reaction, for which we used annotations for the subunits of mitochondrial ribosomal subunits from (Greber & Ban, 2016; Woellhaf et al., 2014).

**Proteome Maintenance Constraint 1: Folding of the proteins**

Chaperone-mediated folding was considered for the cytosolic proteins by multiple chaperone systems. We acquired the information on the clients of these chaperones from the chaperon atlas (Gong et al., 2009). For the cytosolic proteins without a chaperone specified, we assumed folding by chaperone SSB, one of the most abundant and well-studied chaperones. All the proteins in the mitochondria were assumed to be folded by mitochondrial isoform (mt) of Hsp70.

Thus, for each chaperone, the flux through its complex formation reaction could be computed from the sum of the fluxes through protein folding reactions for all its clients:

$$\sum_{e_i} \frac{1}{m} v_{folding,i} \leq k_{cat,chaperone} \frac{v_{complex\, formation,chaperone}}{\mu + k_{deg,chaperone}}$$

where $m$ is a number of chaperones, known to fold the same client and $k_{cat,chaperone} = 0.95\, min^{-1}$ is a turnover constant, reported from ATP turnover measurements for SSB (Lopez-Buesa et al., 1998). Turnover values of the same order were also obtained for chaperone BiP (Rosam et al., 2018).

##### Proteome Maintenance Constraint 2: Degradation of the proteins

For most of the model proteins, protein degradation is described as a proteasome-dependent process. However, the 8 proteins produced by mitochondrial ribosome are assumed to be degraded by a mitochondrial protease PIM1 instead. All protein degradation reactions were coupled with the complex formation of the proteases (either proteasome complex or PIM1) using the following constraint:

$$\sum_{e_i} v_{deg,e_i} \leq k_{cat,protease} \frac{v_{complex\, formation,protease}}{\mu + k_{deg,protease}}$$

where $k_{cat,\mathrm{protease}}$ is the catalytic turnover rate of the protease. The proteasome (subunit composition reported in (Finley et al., 2012)) can degrade proteins with a turnover rate of $k_{cat,proteasome} = 2.3\, min^{-1}$ (Peth et al., 2013), while for PIM1, we used the value reported for its homolog in *E. coli*, Lon protease: $k_{cat,Lon} = 0.26\, s^{-1}$ (Patterson-Ward et al., 2007). For both complexes, we set the cost of protein degradation to be 1.35 molecules of ATP per amino acid (Hong et al., 2012).

##### Proteome Maintenance Constraint 3: Protein transport to mitochondria

Proteins are transported to mitochondria by the TOM-TIM translocase system (Fox, 2012). TOM complex is located in the outer mitochondrial membrane, while TIM22 and TIM23 are located in the inner membrane. All the proteins first enter the mitochondrial intermembrane space through TOM. Next, proteins of the inner mitochondrial membrane (IMM) and mitochondrial matrix are transported into the matrix via TIM23. Then, the IMM proteins are inserted from the matrix side by TIM22. Thus we define a set of mitochondrial transporters $MT = \{TOM, TIM22, TIM23\}$ and using the annotations of the localization of these proteins, define which transporters (or combinations of these) are used for which transport processes.

$$\sum_{e_i} v_{transport,e_i,MT_j} \leq k_{cat,MT_j} \frac{v_{complex\, formation,MT_j}}{\mu + k_{deg}}$$

The two main forces controlling protein transport to and out of the mitochondria are (i) membrane potential and (ii) force generation by mitochondrial chaperones, mainly mtHsp70. (Okamoto et al., 2002) reported that a single cycle of mtHsp70 (ATP hydrolysis) creates a power stroke of ca. 3.5 nm (10 residues of the unfolded protein chain), we set additional requirement for ATP hydrolysis in mitochondria:

$$mtHsp70\ ATPase\ (ATP_{mtHsp70}): ATP + H_2O \rightarrow ADP + P_i + H^+$$

$$\sum_{e_i} \frac{l_{e_i}\,[aa]}{10\ [aa\ stroke^{-1}]}\ v_{transport,e_i,TIM23} = ATP_{mtHsp70}.$$

**Proteome Maintenance Constraint 4: Balance of complex degradation and dilution by growth**

As for the metabolic proteins, we define the same balance of complex formation and their dilution by growth for all the macromolecular complexes (ribosomes, translation factors, chaperones, proteases etc.) in the model:

$$v_{degradation} = k_{deg} \frac{v_{complex\ formation}}{\mu + k_{deg}} \text{ and } v_{dilution} = \mu \frac{v_{complex\ formation}}{\mu + k_{deg}}.$$

#### 2.4 Compartment-specific proteome constraints

**Proteome Constraint 1: Total proteome mass constraint**

(Canelas et al., 2011) observed that the sum mass of the proteins in dry cell mass for the CEN.PK 113-7D strain is growth rate-dependent. Notably, in the measurements of (Canelas et al., 2011), the protein content and water content were reported separately, but since water is a byproduct of protein polymerization, we added these two components together. We implemented the following constraint, so that at any given growth rate, the protein content would correspond to the experimentally determined protein content in dry cell biomass $f_{p,biomass}(\mu)$.

$$\sum \frac{v_{syn,i}}{\mu + k_{deg}} [mmol/gDW] \times \frac{MW_i \left[\frac{g\ protein}{mol}\right]}{1000\ \left[\frac{mmol}{mol}\right]} = f_{p,biomass}(\mu)\ \ [gram\ protein/gDW]$$

where $f_{p,biomass}(\mu) = 0.4854\,\mu + 0.3888$.

**Proteome Constraint 2: Cytoplasmic volume constraint**

In order to establish a constraint on the cytoplasmic volume, we assumed that all the peptides $e_i$, which are synthesized by either the cytosolic or mitochondrial ribosome, occupy *total* cytosolic volume regardless of their final compartment of action. For each protein $e_i$, we estimated volume of a single molecule of protein: (Erickson, 2009) assumed that the protein molecule is a sphere and its radius $r$ was assumed to be a function of molecular weight $MW$:

$$r_i = 0.066\, MW_i^{\frac{1}{3}}$$

We then used $r_i$ to estimate the volume and area occupied by each protein:

$$V_i = \frac{4}{3}\pi r_i^3$$

We used the data of (Tyson et al., 1979) to estimate the growth-dependent volume $V_{cell}(\mu)$ of single cells of *S. cerevisiae*, which vary from ca. $V_{cell} = 20.5\ \mu m^3$ (at $\mu \rightarrow 0$) to $V_{cell} = 32.6\ \mu m^3$ (at $\mu = 0.41\ h^{-1}$).

Then we used the data of (Canelas et al., 2011) to estimate the protein-available fraction $f_{pa}(\mu)$ of the total cell volume, so that the sum mass of the proteins would correspond to the protein content in dry cell biomass $f_{p,biomass}(\mu)$, determined for the CEN.PK 113-7D strain by (Canelas et al., 2011).

The protein-available fraction $f_{pa}(\mu)$ can be decomposed into two variables, $f_{pa}(\mu) = c \times f_{p,biomass}(\mu)$, where $c$ is a growth rate-independent conversion factor. We can estimate the value of $c$ by using the following example for growth rate $\mu = 0$. For any growth rate, the following is true (see Proteome Constraint 1):

$$\sum \frac{v_{syn,i}}{\mu + k_{deg}} [mmol/gDW] \times \frac{MW_i \left[\frac{g\ protein}{mol}\right]}{1000 \left[\frac{mmol}{mol}\right]} = f_{p,biomass}(\mu)\ \ [gram\ protein/gDW]$$

Considering a single molecule of protein YDRUMD1 ($MW_{YDRUMD1} = 48870\ g/mol$), we can estimate both volume and mass of a single molecule of YDRUMD1:

$$r_{YDRUMD1} = 0.066 \times (MW_{YDRUMD1})^{\frac{1}{3}}$$

$$V_{YDRUMD1} = \frac{4}{3}\pi r_{YDRUMD1}^3 = 58.85\ nm^3$$

$$m_{YDRUMD1} = MW/N_A = 8.115 \times 10^{-20}\ g$$

At zero growth $\mu = 0$, the $f_{p,biomass}(0) = 0.3888\ g/gDW$. Considering this mass to be composed solely of YDRUMD1 molecules, we can compute the volume these molecules would occupy:

$$V_{prot} = \frac{V_{YDRUMD1} \times f_{p,biomass}(0)}{m_{YDRUMD1}} = 2.820 \times 10^{11} \mu m^3$$

(Canelas et al., 2011) also estimated that the volume of 1 gram of dry cell weight is $V_{gDW} = 1.7\ mL/gDW$, which was reported to remain relatively constant given different growth rates. Thus by computing the ratio of the volume, occupied by YDRUMD1, and the total cell volume, we can yield the $f_{pa}(0)$:

$$f_{pa}(0) = \frac{V_{prot}}{V_{gDW}} = 0.1658$$

Finally, by knowing the $f_{p,biomass}(0)$ and $f_{pa}(0)$, we can compute $c$:

$$c = \frac{f_{pa}(0)}{f_{p,biomass(0)}} = 0.426$$

It should be noted that this computation is valid regardless of just single protein species taken, as the relationship between the molecular weight and the volume of the sphere as described by (Erickson, 2009) is linearly-dependent, thus any protein would fulfill the requirements.

To estimate the contribution of any protein to the protein-available volume, we used the reasoning of (Erickson, 2009) further. For every protein species $e_i$, we calculated the volumes of single protein molecules in $nm^3$, and considered the following relationship to acquire the volume in $\mu m^3/mmol$ of protein:

$$V_{molar,i} = V_i \times 10^{-9} \frac{\mu m^3}{nm^3} \times 6.022 \times 10^{20}\ mmol^{-1}$$

By using the volume of 1 gram of dry cell weight $V_{gDW} = 1.7\ mL/gDW$, we can estimate the number of cells in a gram of cell dry weight:

$$N_{cells} = \frac{V_{gDW} \times 10^{-6} \frac{m^3}{mL}}{V_{cell} \times 10^{-18} \frac{m^3}{\mu m^3}}$$

and then distribute the molar volume to the number of cells in 1 gram of dry cell weight, yielding how much volume would a 1 $mmol/gDW$ of protein occupy in a single cell:

$$V_{i,single\ cell} = \frac{V_{molar,i}}{N_{cells}}$$

Then we formulate the constraint as follows:

$$\sum \frac{v_{syn,i}}{\mu + k_{deg}} \times V_{i,single\ cell} = V_{cell}(\mu) \times f_{pa}(\mu).$$

##### Proteome Constraint 3: Mitochondrial volume constraint

Analogously to the all the total cytosolic volume constraint, we formulated a mitochondrial protein volume constraint, which constrains the sum volume of all model proteins allocated to this compartment (including mitochondrial ribosomes and import machinery). We formulated the following constraint:

$$\sum_{j \in Mito} \frac{v_{syn,j}}{\mu + k_{deg}} \times V_{j,single\ cell} = V_{cell}(\mu) \times f_{pa}(\mu) \times f_{ma}$$

The mitochondrial proteome volume is always expressed as a fixed fraction $f_{ma}$ (for "mitochondrial available") of the total volume available to the proteins $f_{pa}(\mu)$. We set this fraction $f_{ma} = 0.17$, which sets the available volume of mitochondrial proteins ($c \times f_{ma}$) to about 7.25% of the total protein-available cellular volume, which is close to the 7.4% reported by (Visser et al., 1995).

##### Proteome Constraint 4: Inner mitochondrial membrane constraint

Furthermore, all inner mitochondrial membrane (IMM) proteins from the metabolic model (e.g. electron transport chain proteins, protein translocation complexes and transporters for metabolites) were used to formulate a constraint on IMM surface area. Since the defining factor in this case is the area of the cross-section of these proteins, which spans the membrane (and for globular proteins is defined as $A_i = \pi r_i^2$), we modified the previously derived relation to account for the IMM surface area:

$$A_{molar,i} = A_i \times 10^{-6} \frac{\mu m^2}{nm^2} \times 6.022 \times 10^{20}\ mmol^{-1}$$

and then distribute the molar area to the number of cells in 1 gram of dry cell weight (see Proteome Constraint 1), yielding how much IMM surface area would a 1 $mmol/gDW$ of protein occupy in a single cell:

$$A_{i,single\ cell} = \frac{A_{molar,i}}{N_{cells}}$$

And formulated the constraint as follows:

$$\sum_{j \in IMM} \frac{v_{syn,j}}{\mu + k_{deg}} \times A_{j,single\ cell} \leq A_{IMM}$$

(Lynch & Marinov, 2017) and (Uchida et al., 2011) reported that the surface area of the mitochondrial inner membrane is about 15 $\mu m^2$ in rich YPD medium. We used these measurements to estimate of the maximal mitochondrial membrane area.

Compared to $D = 0.27\ h^{-1}$ in glucose limited chemostats, where maximal mitochondrial protein expression is the highest, the growth rate on YPD-glucose is about $0.45\ h^{-1}$, and $0.4\ h^{-1}$ on minimal glucose medium. Therefore, if we extrapolate the drop in mitochondrial proteins (Figure 4d in the main text), then the mitochondrial content on YPD must be lower than on minimal media in batch. As shown by (Di Bartolomeo et al., 2020), the fraction of the cell volume, occupied by mitochondria, can differ up to 6-fold between cells grown in glucose-rich medium (low number of mitochondria) and ethanol-growing (high number of mitochondria) cells. Then the 6 times decrease in mitochondrial protein content on YPD reported in literature is consistent with our proteomics data that show that maximal mitochondrial protein expression (glucose-limited chemostats, $D = 0.27\ h^{-1}$) is about 4 times higher than on glucose minimal media in batch ($\mu = 0.4\ h^{-1}$).

Assuming that mitochondrial inner membrane area-to-volume ratio is constant, in this extreme case, a 6-fold increase in volume fraction would correspond to a 6-fold increase in mitochondrial inner membrane surface area, setting the upper bound to ca. 90 $\mu m^2$. As (Krauss, 2001) reported that proteins occupy about 80% of the mitochondrial inner membrane area, we can account for that by reducing the upper bound by 20%, to 72 $\mu m^2$. In the end, we set the upper bound of surface area of the inner mitochondrial membrane to 75 $\mu m^2$ in our model.

##### Proteome Constraint 5: Plasma membrane constraint

We previously used data from (Tyson et al., 1979) to determine the volume of a single cell. Assuming a cell of *S. cerevisiae* to be sphere-shaped, we can estimate the plasma membrane surface area using the volume of the cell:

$$A_{PM} = 4 \times \pi \times r^2$$

where $= \left(\frac{3}{4\pi} V_{cell}\right)^{\frac{1}{3}}$ , a radius of a spherical cell.

To estimate the surface area the cross-section of these proteins occupy, we used the previously derived relation (Proteome Constraint 4) to account for the plasma membrane surface area:

$$A_{molar,i} = A_i \times 10^{-6} \frac{\mu m^2}{nm^2} \times 6.022 \times 10^{20}\ mmol^{-1}$$

and then distribute the molar area to the number of cells in 1 gram of dry cell weight (see Proteome Constraint 1), yielding how much PM surface area would a 1 $mmol/gDW$ of protein occupy in a single cell:

$$A_{i,single\ cell} = \frac{A_{molar,i}}{N_{cells}}$$

And formulated the constraint as follows:

$$\sum_{j \in PM} \frac{v_{syn,j}}{\mu + k_{deg}} \times A_{j,single\ cell} \leq A_{PM}.$$

#### Proteome Constraint 6: Glucose transport membrane area

During model development, it became clear that at low saturation of the glucose transporter, mimicking low dilution rates of glucose-limited chemostat cultures, we overestimated growth rate due to all the membrane being predicted to be occupied with glucose transporters. In chemostat cultures, (VanKuyk et al., 2004) measured glucose transport kinetics at different dilution rates and concluded that the maximal rate of glucose transport is rather constant, with a $v_{max} = 620\ nmol/min/mg\ protein$ , equivalent to $v_{max} = 620 \times 10^{-6} \times 60 \times 500 = 18.6\ mmol/gDW/h$, which is close to the maximal glucose consumption rate of approximately 16 to 19 $mmol/gDW/h$, documented by (Blank & Sauer, 2004). With an average $k_{cat} = 125\ s^{-1}$ of the various glucose transporters, taking the flux interval reported by (Blank & Sauer, 2004) as the upper bound, we compute that glucose transport covers about 7.5 $\mu m^2$. We used this number as upper limit:

$$A\ ([HXT]) \leq 7.5\ \mu m^2$$

This is an empirical proteome constraint, not based on fundamental limits but on regulatory mechanisms. The fact that it is an active constraint in wild-type yeast is confirmed by laboratory evolution experiments in glucose limited chemostats, which have consistently showed increases in high-affinity glucose transporter expression (Ferea et al., 1999).

#### Proteome Constraint 7: Unspecified protein

The model does not include a complete list of proteins that might be expressed in the organism: only proteins known to catalyze metabolic reactions are included in genome-scale metabolic models. Thus we need to account for the synthesis of proteins required for storage, signaling or maintaining the structure of the cell. This is especially important at low growth rates, where only low amounts of enzymes are needed to sustain the low metabolic fluxes.

In the biomass reaction, the composition of amino acids is given with stoichiometric coefficients in $mmol/gDW$. We multiplied these coefficients with the molecular weight (MW) of each of the amino acids ($g/mmol$), to convert the amino acid composition $r$ as $mmol\ aa/mmol$ of total amino acids. As a proxy for the proteins we do not explicitly account for, and assuming the length of an average protein to be around 450 amino acids, we designed a dummy protein YDRUMD1. Therefore, the number of residues of a certain amino acid in the unspecified protein YDRUMD1 chain equals to $450 \times r$. Additionally, we set that YDRUMD1 has 50 nucleotides in its 5'UTR, the median for other proteins.

Even in high growth rates, this *unspecified protein* ($UP$) should be still expressed to a certain degree, as otherwise all the proteome would be occupied by metabolic enzymes – being a rather unrealistic scenario. Using the proteomics data, we estimated about $0.25\ g/g\ Prot$ of proteins. This decision was based on the comparison of proteome fractions, allocated to different cellular processes ($g/g\ total\ protein$), from experimental data and model predictions (Fig. S1).

For this, we attributed proteins to coarse proteome sectors. The first category of proteins was associated with proteome turnover (similar to Terry Hwa's "R" fraction): translation factors, ribosomal proteins, chaperones, proteasome subunits/proteases (in all cases, including not only cytoplasmic, but also their mitochondrial counterparts). The remaining proteins from experimental datasets were attributed to another category ("Rest of cellular processes", a combination of Hwa's "P" and "Q" sectors). In the case of experimental data (points in the Fig. S1), these proteins comprised not only metabolic proteins (shown as orange triangles in the plot) but also proteins with other functions (such as signaling and structural proteins).

Then we compared the experimental measurements with model simulations, where the lines show either, again, the proteome fraction of proteins, involved in protein turnover, or - for the "Rest of cellular processes" - the sum of proteome fractions occupied by metabolic proteins and the UP. Notably, with the minimal UP value of $0.25\ g\ UP/g\ protein$, we do see an agreement of the model predictions to the experimental data for both the protein fractions, assigned to both "Proteome turnover" and "Rest of cellular processes" categories. Therefore, we believe that the data supports our decision to use the UP value of $0.25\ g\ UP/g\ protein$ for model simulations.

Thus, we set the minimal requirement of the $UP$ in the proteome to $0.25\ g/g\ Prot$:

$$\frac{v_{syn,UP}}{\mu + k_{deg}}[mmol/gDW] \times \frac{MW_U[g/mol]}{1000\ [mmol/mol]} \geq UP\ [g/g\ prot]\ \times\ prot\ ratio\ [g\ prot/gDW]$$

Subsequently, the balance between the degradation and dilution of the *UP* is defined as follows:

$$v_{deg,U} = k_{deg,U}\frac{v_{syn,U}}{\mu+k_{deg,U}} \text{ and } v_{dil,U} = \mu\frac{v_{syn,U}}{\mu+k_{deg,U}}.$$

#### Proteome Constraint 8: Gratuitous protein expression

We included descriptions of some gratuitous proteins (GFP, YFP, mCherry and haemoglobin) into the model to simulate stress response to gratuitous protein expression. As for other proteins, we added protein translation, folding, degradation and dilution-by-growth reactions. The fraction of the total proteome that the gratuitous protein occupies is described by the following constraint (taking $mCherry$ as the example):

$$\frac{v_{syn,mCherry}}{\mu + k_{deg}}\left[\frac{mmol}{gDW}\right] \times \frac{MW_{mCherry}\left[\frac{g}{mol}\right]}{1000\ \left[\frac{mmol}{mol}\right]} = mCherry\ [\frac{g}{g\ prot}]\ \times\ prot\ ratio\ [\frac{g\ prot}{gDW}]$$

Similarly to the UP, the balance between the degradation and dilution of the $mCherry$ is defined as follows:

$$v_{deg,mCherry} = k_{deg,mCherry}\frac{v_{syn,mCherry}}{\mu+k_{deg,mCherry}} \text{ and } v_{dil,mCherry} = \mu\frac{v_{syn,mCherry}}{\mu+k_{deg,mCherry}}.$$

Inhibition of growth (reduction in maximal achievable growth rate) by gratuitous protein expression is simulated by pre-setting the fraction of proteome which gratuitous protein occupies, and running the binary search in order to find the highest growth rate, still resulting in a feasible linear program (just as for unperturbed state).

#### 2.5 Growth-related constraints

##### Growth-related Constraint 1: Biomass equation

The biomass reaction describes the amount of biomass precursors as $mmol/gDW$ (Thiele & Palsson, 2010). (Forster et al., 2003) described how the concentrations these precursors were computed from published data. The composition of carbohydrates, proteins (as amino acids), DNA, and RNA (as [deoxy]ribonucelotides) is included to the biomass equation.

In the biomass equations of the conventional GEMs, the stoichiometric coefficients of the precursors do not change with the growth rate. However, the biomass composition is known to vary from strain to strain and, in the case of CEN.PK113-7D, was observed to be a function of growth rate (Canelas et al., 2011). Thus, we used the data of (Canelas et al., 2011) to obtain the regression curves for the growth rate-dependent change in bulk biomass composition, when the growth rate is determined by different glucose availability (SN Table 1). Thus, for simulating glucose-limited growth, at every simulation step, we set the flux through the new biomass reaction to be equal to the corresponding growth rate and recompute its composition according to the relationships listed in SN Table 1.

However, it is known that cells, treated with the translation inhibitor cycloheximide do not accumulate storage carbohydrates despite the halted growth (Thomsson et al., 2005). Thus for simulating inhibition of ribosomes, we set the biomass composition to be that of $\mu = 0.40\ h^{-1}$ at all times. Notably, we also performed analogous ribosome inhibition simulations where the biomass composition was changing depending on the growth rate $\mu$ as described above, however, that did not have a major effect to the outcome of the simulations.

**SN Table 1**. Dependency of the abundance of biomass precursors on the growth rate changed by glucose availability.

| Component | Regression equation (or fixed amount) $f = [g/gDW]$ |
|---|---|
| Protein | $f_{p,biomass} = 0.4854\,\mu + 0.3888$ |
| Lipids | $f_{lipid} = -0.0456\,\mu + 0.0745$ |
| RNA | $f_{RNA} = 0.1525 \times \mu + 0.0498$ |
| Adenylate* | $f_{Ade} = 0.2 \times f_{RNA}$ |
| Guanylate* | $f_{Gua} = 0.283 \times f_{RNA}$ |
| Cytidylate* | $f_{Cyt} = 0.26 \times f_{RNA}$ |
| Uridylate* | $f_{Ura} = 0.20 \times f_{RNA}$ |

| | |
|---|---|
| DNA** | $f_{DNA} = 0.005$ |
| Inorganic phosphate | $f_{P_i} = 0.006$ |
| Inorganic sulphate | $f_{SO_4^{2-}} = 0.003$ |
| Carbohydrates (total)*** | $f_{carbs} = -0.5249 \times \mu + \ 0.449$ |
| Trehalose**** | $f_{treh} = (-0.555 \times \mu + 0.183) \times f_{carbs}$ |
| Glycogen**** | $f_{glyc} = (-0.837 \times \mu + 0.317) \times f_{carbs}$ |
| Mannans | $f_{treh} = (0.511 \times \mu + 0.219) \times f_{carbs}$ |
| Glucans | $f_{treh} = (0.880 \times \mu + 0.281) \times f_{carbs}$ |

* - data from (Tada & Tada, 1962);
** - exact composition of the deoxyribonucleotides taken from the Yeast7.6;
*** - data from (Lange & Heijnen, 2001);
**** - if the computed fraction is less than zero, the value becomes zero;

To define the growth rate-dependent lipid composition, we made a lumped reaction consisting of 64 lipid species. We used the data of (Ejsing et al., 2009), where the data were provided for every lipid species as mol/mol lipid. First, we recomputed the molar fractions into mass fractions, using the molecular masses of individual lipids from the Yeast Metabolome Database (YMDB) (Ramirez-Gaona et al., 2016) and LIPID MAPS (http://www.lipidmaps.org/). Then, using the bulk measurements of (Canelas et al., 2011), we distributed the total lipid mass fraction to the individual lipids, including them into the biomass equation in $[mmol/gDW]$.

In the case of inhibition of translation at glucose excess, we kept the biomass composition that corresponds to the uninhibited state.

##### Growth-related Constraint 2: ATP maintenance

For the growth-related processes that we do not account for explicitly in the model, we need to define an ATP hydrolysis reactions, that are split into a growth-associated ATP maintenance, GAM, and a non-growth associated maintenance reaction NGAM:

$$GAM: ATP + H_2O \rightarrow ADP + P_i + H^+$$
$$NGAM: ATP + H_2O \rightarrow ADP + P_i + H^+$$

The GAM value is multiplied by the growth rate ($v_{GAM} \times \mu$), whereas NGAM is fixed to the NGAM value $v_{NGAM}$. For galactose and maltose growth, the added a third ATP maintenance term cNGAM to test if additional ATP cost would improve the predictions (see Section 5 of this document).

In conventional genome-scale metabolic models, e.g. Yeast7.6, the flux of GAM $v_{GAM}$ at the growth rate $\mu^*$ is defined as $v_{GAM}(\mu) = 59.3\, mmol/gDW \ \times \mu$. However, our model already describes many of the ATP-investing processes, as ATP is used in protein translation, folding and degradation. As reported by (Lahtvee et al., 2017), at $\mu = 0.1\ h^{-1}$, the protein biosynthesis and turnover costs are about 23 and 12 $mmol/gDW$, respectively.

Experimentally, the value of GAM is usually estimated using flux data from glucose-limited chemostats (Verduyn et al., 1990b, 1990a), and, in the case of aerobic growth, always from fully respiratory data below the critical dilution rate. Following these procedures, we found that a GAM value of $v_{GAM}^{max}(\mu) = 24.0\ mmol/gDW \times \mu$ was able to describe full respiratory growth on glucose very well (Fig. 2 in the main text). However, our initial simulations showed that this value does not fit the data at high growth rates very well: fixing the GAM value resulted in elevated demand of carbon source (in this case, glucose), as well as ca. 30% higher ethanol flux at $\mu \to 0.4\ h^{-1}$ than experimentally reported (both including our own data and that of (Van Hoek et al., 1998)). Thus we concluded, based on the available data, that GAM should be lower at higher growth rates to explain the growth-rate associated fluxes. We have no mechanistic explanation, but since the deviation was the strongest at the highest growth rates, we suspect it to be linked to glucose-dependent signaling. We therefore coupled the GAM value to the saturation level of the hexose transporters $f_{sat}^{HXT}$:

$$v_{GAM}(\mu, f_{sat}^{HXT}) = v_{GAM}^{max} \times \mu \times (1 - 0.5 f_{sat}^{HXT})$$

As the saturation of the hexose transporters $f_{sat}^{HXT}$ is low, the $v_{GAM} \approx 24.0\ mmol/gDW \times \mu$; low saturation rates ($f_{sat}^{HXT} < 0.20$) indeed correspond to the glucose-limited growth with $D < D_{crit}$, keeping the consistency with the experimental observations. As $f_{sat}^{HXT} \to 1$, $v_{GAM} \to 0.5\ v_{GAM}^{max} \times \mu$, or $v_{GAM} \approx 12\ mmol/gDW \times \mu$.

To estimate the value of the NGAM, we used the flux data from glucose-limited conditions: aerobic cultivation in chemostat by (Canelas et al., 2011; Van Hoek et al., 1998) and this study, as well as cultivation in anaerobic chemostats (Nissen et al., 1997). Plotting the $v_{ATP}(\mu)$, we found an NGAM value of $v_{ATP}(0) = v_{NGAM} = 0.818\ mmol/gDW/h$ (FigSN1). However, this ATP expenditure still included the costs of synthesis and degradation of the proteins at near-zero growth. To account to this, we computed the respective requirements of ATP ($v_{ATP}^{p-syn}(0) = 0.486\ mmol/gDW/h$ and $v_{ATP}^{p-deg}(0) = 0.164\ mmol/gDW/h$) and, by subtracting these components from the total NGAM, we set the $v_{NGAM} = 0.168\ mmol/gDW/h$ in our model.

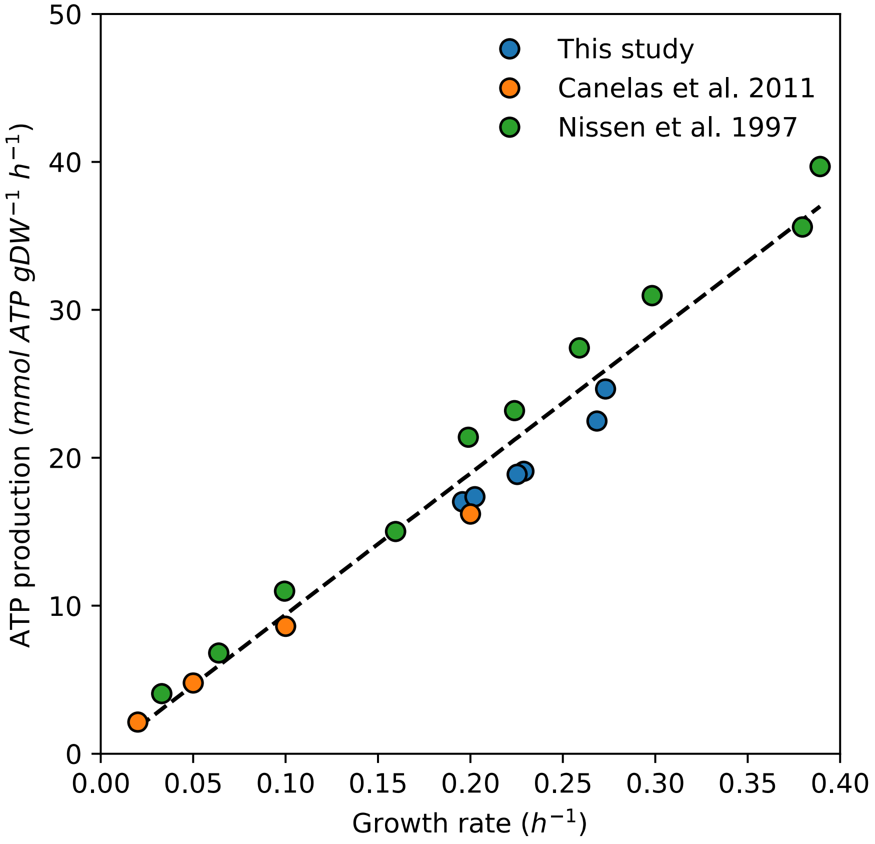

**Fig. SN1.** Computed ATP production for different growth rates in anaerobic (Nissen et al., 1997) and aerobic ((Canelas et al., 2011), this study) settings. ATP production was computed as $v_{ATP}^{aerobic} = 2 \times v_{Glc} + \frac{1}{3} \times v_{CO_2} + 2 \times v_{O_2}$ for aerobic conditions (representing the production by glycolysis, TCA cycle and oxidative phosphorylation, respectively) and $v_{ATP}^{anaerobic} = 2 \times v_{Glc} - v_{Glyc}$ (production by glycolysis, corrected for carbon, rerouted to glycerol).

##### Growth-related Constraint 3: Growth media composition (exchange reactions)

The minimal growth medium we considered contained glucose, sulphate, ammonia, and phosphate. To mimic different growth media, we changed the upper and lower bounds for exchange reactions of the metabolites, according to the descriptions of the SC and YPD growth media from (Snitkin et al., 2008). For the minimum glucose medium, we set both lower and upper flux bounds to zero for all exchange reactions for amino acids and other metabolites, which are otherwise present in rich growth media. For simulating anaerobic growth, the minimal glucose-based media was also supplemented with trace amounts of ergosta-5,7,22,24(28)-tetraen-3beta-ol (an ergosterol derivative, crucial for growth), palmitoleate and oleate (the latter two are experimentally provided with addition of Tween 80 to the growth medium).

In most of the experiments we performed ourselves and collected from literature sources, glucose was used as the sole (or main, in case of rich SC/YPD media) carbon source. Thus, assessment of residual glucose levels are an important consideration on correctly predicting metabolic behaviour. We computed the concentration of residual glucose from the model simulations, using formula $[Glc_{Res}] = \frac{f_{sat}}{1-f_{sat}} \times K_M$, with $f_{sat}$ being the saturation factor of the glucose transporters and $K_M = 1\, mM$ being the Michaelis constant of high-affinity glucose transporters. Predictions of the residual glucose levels agree to the experimental observations that suggest only minute amounts of glucose in spent media from glucose-limited chemostats (see Fig. 2A of the main text).

We modeled glucose uptake as irreversible process and, as a result, obtained good agreement with the experimental data on both the residual glucose concentrations and glucose uptake rates. Therefore, we conclude that we can safely neglect the potential inhibitory effects of intracellular glucose concentration. In glucose-limited chemostats, the concentration of intracellular glucose is expected to be very low and the effect therefore very small. It should be noted that a strong impact of intracellular glucose would only complicate the mapping between saturation level of the transporter and the extracellular glucose concentration, but not the effect of transport saturation on growth rate and metabolic strategies.

**Growth-related Constraint 4: Energetics of oxidative phosphorylation**

Measurements of ATP yields from oxidative phosphorylation in S. cerevisiae show that this process is far away of its theoretical yield of ca. 32 $ATP/Glc\ [mol/mol]$. Rather than that, usually reported value is twice as small, ca. 16 $ATP/Glc$ (Bakker et al., 2001). However, in the original Yeast7.6 model, maximal yield of ATP is $22\, mol\ ATP/mol\ Glc$ ($\max ATPase$, $v_{Glc} = 1\, mmol/gDW/h$, exchange for $O_2$, and inorganic ions unbound).

(Bakker et al., 2001) argued that one of the main factors leading to low yield of ATP from oxidative phosphorylation is proton leaks through the inner mitochondrial membrane from the intermembrane space towards the matrix. This phenomenon was observed already decades ago (Garlid et al., 1989), was shown to be dependent on the intensity of respiration (Fitton et al., 1994) and was suggested to be an active process (Rigoulet et al., 2010).

To account for this process, we implemented a linear constraint, which forces two protons to leak from intermembrane space to the mitochondrial matrix for every pair of electrons transferred from NADH onto ubiqui-6-one. In Yeast7.6, such constraint changes the ATP yield to 16 $mol\ ATP/mol\ Glc$.

### 3. Manual curation of turnover values

#### 3.1 Maltose-specific metabolism

Compared to glucose, to shuttle maltose into glycolysis, two new enzymes have to be expressed: a maltose-specific transporter MAL11 and maltase MAL12. Even though we collected a high number of turnover values, there is still high uncertainty about many of these enzyme turnover values, which are directly linked to the protein costs in our model. We noticed inconsistencies between the retrieved turnover values and experimental data for maltase: based on experimental data, in maltose-grown cells, enzyme maltase makes up about 2% of the total proteome. Meanwhile, the reported turnover value was $k_{cat}^{GECKO} = 1278.0\ \mathrm{s}^{-1}$, which is unrealistically high given the measured proteome fraction. Therefore, we manually adjusted the turnover values of the maltase using our proteomics data.

We first obtained values for $v_{max}$ for individual enzymes from reports in the literature. We then computed the amount of enzyme in the dry mass of *S. cerevisiae* $[e_i]$ from our proteomics data, using the relationship between the growth rate and protein content in dry mass from (Canelas et al., 2011) as follows: by knowing the proteome fraction in $g/g\ protein$ and the total amount of proteins in dry biomass ($g\ protein/gDW$), we computed the mass of protein of interest $m_i$ (in $g/gDW$), which we converted into amount of protein $[e_i]$ (in $mmol/gDW$) using its molecular weight. Assuming that the enzymes of interest are bottlenecks in their respective pathways and thus are fully saturated, we followed the relationship $v_{max} = k_{app} \times [e_i]$, and computed the $k_{app}$ values using the maximal flux values $v_{max}$ we collected (SN Table 2).

We also compared the expression levels and measured turnover values for galactose-specific proteins (the Leloir pathway). Yet, we did not notice striking differences and thus did not adjust the turnover values.

**SN Table 2.** Estimated $k_{eff}$ value for maltase.

| Gene | Protein | Functionality | Proteome mass fraction | $k_{cat}$ (GECKO) ($s^{-1}$) | $k_{eff}$ ($s^{-1}$) | $k_{cat}$ $/k_{eff}$ | Source |
|---|---|---|---|---|---|---|---|
| *MAL12* | YGR292W | Maltase | 0.0201 | 1278.0 | 2.258 | 565.99 | doi |

#### 3.2 Other metabolic enzymes

For collection of the $k_{cat}$ values for metabolic enzymes, we used the GECKO toolbox (Sánchez et al., 2017). For the proteins without a GECKO-provided $k_{cat}$ value, we assumed a default of $k_{cat} = 90\ s^{-1}$. However, during the process of automatic compilation of the turnover values, some non-physiological measurements are acquired (e.g. turnover number of a mutant protein, or a protein with the same function from a very distant species). Thus we also performed manual curation of the $k_{cat}$ values for the following enzymes (SN Table 3, SN Table 4).

**SN Table 3.** Manually curated turnover values ($k_{cat}$ [cur.]) for metabolic enzymes.

| Gene | Protein | Functionality | $k_{cat}$ (GECKO) ($s^{-1}$) | $k_{cat}$ (cur.) ($s^{-1}$) | Source |
|---|---|---|---|---|---|
| *PMA1* | YGL008C | $H^+$-ATPase | 56.8 | 36.0 | doi |
| | | ATP synthase | 20.0 | 120.0 | doi |
| *GLT1* | YDL171C | Glutamate synthase | 70.0 | 52.7 | doi |
| *GLN1* | YPR035W | Glutamine synthase | 52.7 | 7.02 | |
| *GDH3* | YAL062W | $NADP^+$-dependent GDH | 2682 | 13.0 | doi |

| | | | | | |
|---|---|---|---|---|---|
| *GDH1* | YOR375C | | | | |
| *AQY2* | YLL052C | Aquaporin | 90.0 | 900.0 | Unrealistic model predictions (futile cycle) |
| *DLD2*<br>*CYC7* | YDL178W<br>YEL039C | D-lactate dehydrogenase | 1500.0 | 81.0 | Unrealistic model predictions (artificial pathway) |
| *GCV3*<br>*GSD1*<br>*LPD1*<br>*GSD2* | YAL044C<br>YDR019C<br>YFL018C<br>YMR189W | Glycine cleavage system | 1500.0 | 1.0 | Unrealistic model predictions (futile cycle) |
| *SMH1* | YBR263W | Serine hydroxymethyltransferase | 640.0 | 2.0 | doi |

**SN Table 4.** Manually curated plasma membrane transporters.

| Metabolite | Comment | kcat ($s^{-1}$) |
|---|---|---|
| Glucose | There are many different glucose transporters with different $k_{cat}$ and $K_M$ values. (Ye et al., 2001) and (Kruckeberg et al., 1999) estimated that the turnover constants for glucose transporters HXT7 and HXT2 are equal to 200 and 53 $s^{-1}$, respectively. Rather than specifying the $K_M$values, we used a varying saturation level to change glucose availability in simulations, and used an average $k_{cat}$ of glucose transporters $k_{cat,Glc} = 125\ s^{-1}$. | 125 |
| Phosphate | (Shokrollahzadeh et al., 2007) reported that the phosphate transporter Pho84 has the specific activity as 6 $\mu mol/gDW/min$. The concentration of Pho84 is about 300,000 molecules/cell (Chemical et al., 2003), so we can estimate the $k_{cat}$ of Pho84 as: $$k_{cat} = \frac{\frac{6.1\ mmol}{1000\ gDW\ min}}{\frac{300{,}000}{13 * 6.023 \times 10^8}\ \frac{mmol}{gDW}} = 159.2\ min^{-1} \approx 3\ s^{-1}$$ | 3 |
| Ammonium | (Wacker et al., 2014) measured the rate of transported ammonium as 30 to 300 ions per second. | 30 |
| Sulphate | Default value for proteins without GECKO-provided turnover number | 90 |

### 4. Identification of active Elementary Flux Modes

In the Main Text (see "Changes in metabolic strategies are the result of proteome constraints"), we analyzed the model to understand why the metabolic fluxes changed with glucose availability. For this we made use of earlier developed mathematical theory about the solutions of growth rate maximization that use rates as optimization variables and weighted sums of proteins as constraints (de Groot et al., 2019; Wortel et al., 2014). Optimal solutions of such maximization problems are superpositions of so-called Elementary Growth Modes (EGMs), whose metabolic subnetwork can be described by long-known Elementary Flux Modes (EFMs) under constant amino acid composition of the biomass (de Groot et al., 2020).

EFMs can be seen as metabolic pathways with fixed stoichiometry between its substrates and products, and they are elementary in the sense that each reaction in an EFM is essential to maintain steady state (Wortel et al., 2014). It was mathematically proven that the number of EFMs that can carry flux in optimal solutions is bounded by the number of constraints that actively limit the maximal growth rate (de Groot et al., 2019). Which specific flux modes, and hence metabolic strategies, are optimal, depends on the cost-effectiveness (or "proteome efficiency" (Basan et al., 2015; Chen & Nielsen, 2019)) of each mode.

The detailed mapping of all processes in protein synthesis and degradation allows pcYeast to compute the costs of implementing these strategies. In this section, we present the procedure to identify the number of active Elementary Flux Modes (EFMs) in (proteome-constrained) genome-scale models, stressing particularities of this procedure that are specific for the pcYeast model.

As a side note, we first have to discriminate between two types of constraints that we apply to the model: equality constraints and *in*equality constraints. Equality constraints have to be always met for any optimization, while inequality constraints can obtain any values bound by both flux definitions and the right-hand-side. For an inequality constraint on fluxes $v$ with scaling factors $w$: $\sum w_i v_i \leq V$, with all $v_i \geq 0$ and $V \geq 0$, the value of the left-hand-side post-optimization can obtain values in the range of $[0; V]$. We call such an inequality constraint "active" if $\sum w_i v_i = V$ post-optimization.

Following the method initially presented by (Schuster et al., 1999), we begin the enumeration of active EFMs with optimizing the linear program which results in a flux distribution with all non-zero flux values. It should be noted here that all fluxes should be <u>non-negative</u> $v_i \geq 0$ (i.e. reversible reactions are split into two irreversible reactions) and in the pcYeast model, all reversible reactions are split into two unidirectional reactions already (see **Mass Balance Constraint 1**). Note that this list of non-zero fluxes, which we call "active fluxes" will contain either metabolic fluxes alone (conventional genome-scale models) or both metabolic and proteome-related fluxes (in the case of pcYeast). Here we aim to enumerate EFMs and not EGMs (de Groot et al., 2020), and therefore we will consider only the active metabolic fluxes in pcYeast.

We first need to obtain the stoichiometric matrix; we can use the model handling tools, like PySCeS CBMPy (Olivier et al., 2005), to generate the stoichiometric matrix of the model and export it to a text file, which can be handled independently of the software used to run models. Note that for the pcYeast model, we first need to compute the growth rate-dependent stoichiometric coefficients for biomass equation and add the reaction itself, only then export the matrix. The resulting matrix will have a shape of $m$ rows, representing metabolites, and $n$ columns, representing reactions. Having both the list of active metabolic fluxes and the stoichiometric matrix, we can subset the stoichiometric matrix to include only the active fluxes, which we call "active stoichiometric matrix".

We can compute the number of active EFMs by computing the rank of the active stoichiometric matrix or by enumerating EFMs using software packages, like *efmtool* (Terzer & Stelling, 2008). By using the rank computation approach, we compute the rank of the active stoichiometric matrix. The difference between the rank (the number of independent mass balances, i.e. equations) and the number of reactions (the variables) determines the degrees of freedom. Since EFMs have by definition only one degree of freedom, and every flux distribution is a linear combination of EFMs (Schuster et al., 1999), the number of degrees of freedom must equal the number of active EFMs. We have previously reported that such an approach works for conventional genome-scale models of yeast that do not contain equality constraints (Bruggeman et al., 2020).

Applying the rank computation routine is not entirely trivial for the pcYeast model. We performed a computation described above for a glucose-limited condition at low growth rate, where we were expecting a single EFM to be active, based on the observations of number of active compartment-specific proteome constraints, together with stable biomass yields of $Y_{X/S} \approx 0.52\ g/g\ Glc$ and exchange fluxes scaling uniformly with the growth rate (see Fig. 2 in the Main Text). We obtained the difference between the number of active fluxes and the rank of $n - 6$, suggesting 6 EFMs to be active. This result was puzzling at first, but we can show that our claim for the number of growth-supporting EFMs equaling 1 is in fact correct, by taking into account the equality constraints.

In pcYeast we set a number of equality constraints, which represent 3 types of ATP maintenance ($NGAM$, $GAM$, mitochondrial maintenance) and another constraint, representing the proton leak in the mitochondrial inner membrane (see **Growth-related Constraint 3** and **4**, **Proteome Maintenance Constraint 3**). Since the fluxes through the corresponding reactions are either determined prior to optimization (when constraint expression is $v_1 = value$) or linked to other fluxes in the system (e.g. $v_1 = value \times v_2 > 0$), these fluxes are either known both pre- and post-optimization, or can be computed post-optimization from other fluxes. Either way, these fluxes provide additional degrees of freedom in the rank computation.

We have removed each of the equality constraints described above one-by-one, and at each step, the difference between the number of active fluxes and the rank of the corresponding active stoichiometric matrix would be lower by 1. With this, we validated that every additional equality constraint introduces a degree of freedom: with all 4 constraints removed, the rank of the active stoichiometric matrix was $(n - 2)$. The final additional degree of freedom comes from the fact that we fix the value of the biomass flux to the growth rate we simulate at every iteration (**Growth-related Constraint 1**). However, to maintain the steady-state nature of the system, we naturally cannot remove this flux to arrive to the difference of $(n - 1)$.

With some modifications specific to the pcYeast structure we could verify that the result of $(n - 2)$ does correspond to a single EFM, using an alternative approach with the *efmtool*. By running the *efmtool* on the active stoichiometric matrix, the user receives not only the number of EFMs, but also the flux values for every non-zero flux belonging to each of the EFMs. For *efmtool*, we considered that the flux distributions we compute include costs of protein production, both in terms of amino acids and energy. It is important to augment the biomass equation with charged tRNAs as substrates and free tRNAs as products. Moreover, while this is accounted for in the cytoplasmic part of the model, for the sake of completeness, hydrolysis of (even a small amount of) mitochondrial GTP must be included into the biomass equation as well. Subjecting this modified active stoichiometric matrix to the *efmtool* results in detection of one active EFM, containing all the active fluxes with non-zero values.

Thus, by either rank computations or analysis by *efmtool*, we indeed can show that with one compartment-specific proteome constraint active, the predicted flux distribution can be described as a single EFM. The procedure works also well for situations where other proteome constraints are active: for instance, in the case of growth at glucose-excess, the pcYeast suggests only one compartment-specific proteome constraint, minimal UP fraction, to be active. Enumeration of EFMs suggested a single EFM being present, in line with the previously provided example of glucose transporter-limited growth.

### 5. Growth on galactose and maltose

We aimed to simulate batch growth of *S. cerevisiae* on different other hexose sugars, at first with the model that we calibrated with glucose chemostat data without any adjustments ("naïve"), and using the original GECKO-based turnover values for the metabolism of galactose and maltose (except for maltase, see Section 3.1). However, the flux distributions and the growth rates we obtained largely resembled these of glucose excess conditions (SN Table 5), suggesting that the naïve model cannot capture the effects of growth on sugars other than glucose.

**SN Table 5.** Comparison of the experimental measurements and predicted fluxes by the naïve models for growth in excess sugar conditions. Fluxes are in $mmol/gDW/h$ except for the flux through the biomass reaction (the specific growth rate, in $h^{-1}$).

| Substrate | Flux | Experimental data | Naïve prediction |
|---|---|---|---|
| **Glucose** | $\mu\ (h^{-1})$ | 0.390 | 0.400 |
| | $qS_x$ | 13.5 | 13.82 |
| | $q_{O_2}$ | 2.0 | 3.23 |
| | $q_{EtOH}$ | 19.0 | 20.90 |
| | $q_{CO_2}$ | 22.2 | 24.78 |
| **Maltose** | $\mu\ (h^{-1})$ | 0.283 | 0.340 |
| | $qS_x$ | 5.2 | 7.90 |
| | $q_{O_2}$ | 2.7 | 2.62 |
| | $q_{EtOH}$ | 14.1 | 25.96 |
| | $q_{CO_2}$ | 17.0 | 29.15 |
| **Galactose** | $\mu\ (h^{-1})$ | 0.174 | 0.350 |
| | $qS_x$ | 2.6 | 7.61 |
| | $q_{O_2}$ | 5.3 | 5.19 |
| | $q_{EtOH}$ | 0.7 | 8.44 |
| | $q_{CO_2}$ | 6.0 | 14.13 |

We therefore looked for the determinants that could explain the experimentally determined phenotypes of *S. cerevisiae*. First consideration of ours was the maximal uptake of the carbon source: the maximal uptake flux in the naïve model was almost 3-fold and 1.5-fold higher than experimental values for galactose and maltose, respectively. Moreover, the predicted abundance of the galactose- and maltose-specific transporters GAL2 and MAL11, respectively was also between 3-to-10 times higher, compared to quantitative proteomics measurements. Based on these observations, we scanned the maximal area, which hexose transporters can occupy (**Proteome Constraint 6**) (FigSN2) in order to estimate a suitable maximal expression level. Following the results of these simulations, we reduced the maximal area allowed for hexose transporters accordingly: for the galactose case from 7.5 to 3 $\mu m^2/cell$, and for the maltose case from 7.5 to 3.5 $\mu m^2/cell$. It should be noted that we explicitly picked the values which would result in lower uptake fluxes than the experimentally measured.

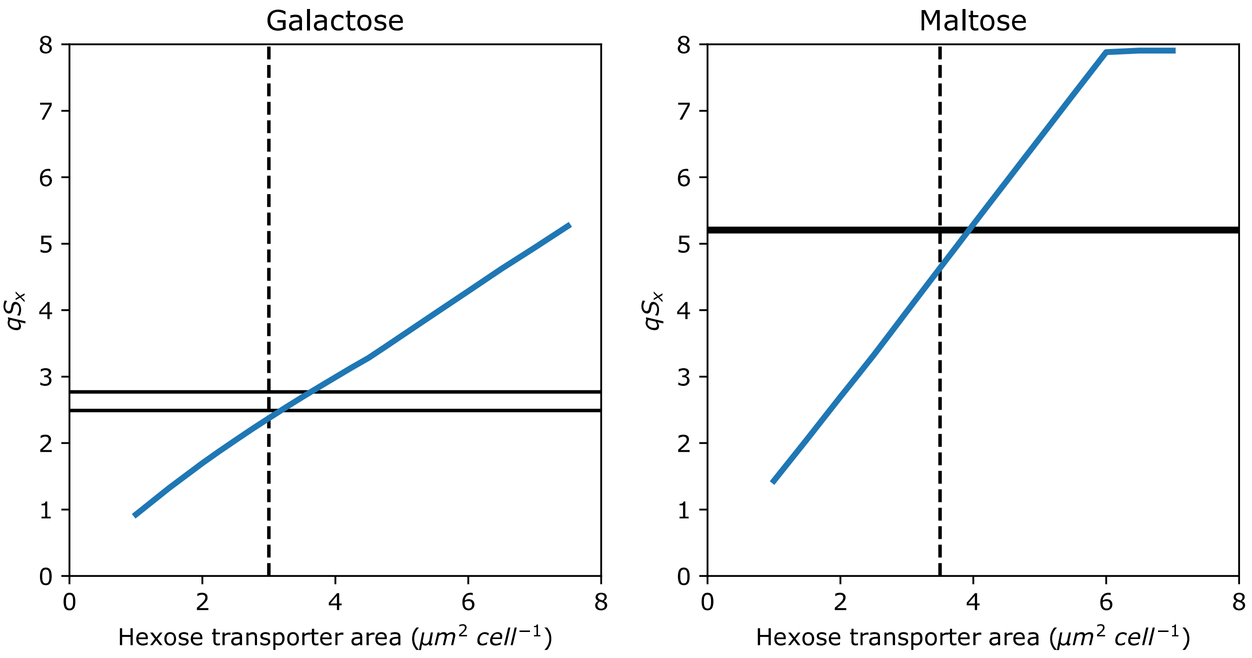

**FigSN2.** Estimation of the maximal area for hexose transporter to occupy. The naïve model was used in simulations with the surface area varying from 1 to 7.5 $\mu m^2/cell$. Solid horizontal lines represent experimentally determined uptake fluxes from individual batch cultures, and dashed vertical lines represent the maximal area which was selected for further experiments.

As we saw a strong negative impact on the experimentally measured growth rate for growth on carbon sources other than glucose, we then postulated that there must be additional effects alongside the apparent expression limit of hexose transporters. Therefore we considered probing how applying additional burden on the total proteome changes the model predictions. This choice was backed by our modeling insight that the growth of *S. cerevisiae* in excess of glucose is limited only by total proteome limitation. For simulations, we used the model and varied the value of minimal unspecified protein (UP) fraction in the proteome in order to impose an additional proteome burden.

Accounting to the initial distance of the naïve predictions from the experimental measurements, we selected to scan different ranges of UP minimal values, between 0.42 and 0.615 $g\ UP/g\ protein$ for galactose. For maltose, we took the same approach but sampled UP minimum between 0.225 and 0.42 $g\ UP/g\ protein$ instead, due to lower reduction of the growth rate, compared to the naïve model. For each of the minimal UP values tested, we acquired the distribution with the highest feasible growth rate. As a criterion of the goodness-of-fit, we then computed the sum of scaled square residuals (compared to the experimental data) based on the flux through the biomass reaction (experimentally determined growth rate), and exchange fluxes of the carbon source, oxygen, ethanol and carbon dioxide: $RSS = \sum\left(\frac{v_i^{exp}-v_i^{pred}}{v_i^{exp}}\right)^2$ (Fig. SN3), meaning that the lower the RSS value, the closer the prediction is to the experimental measurements.

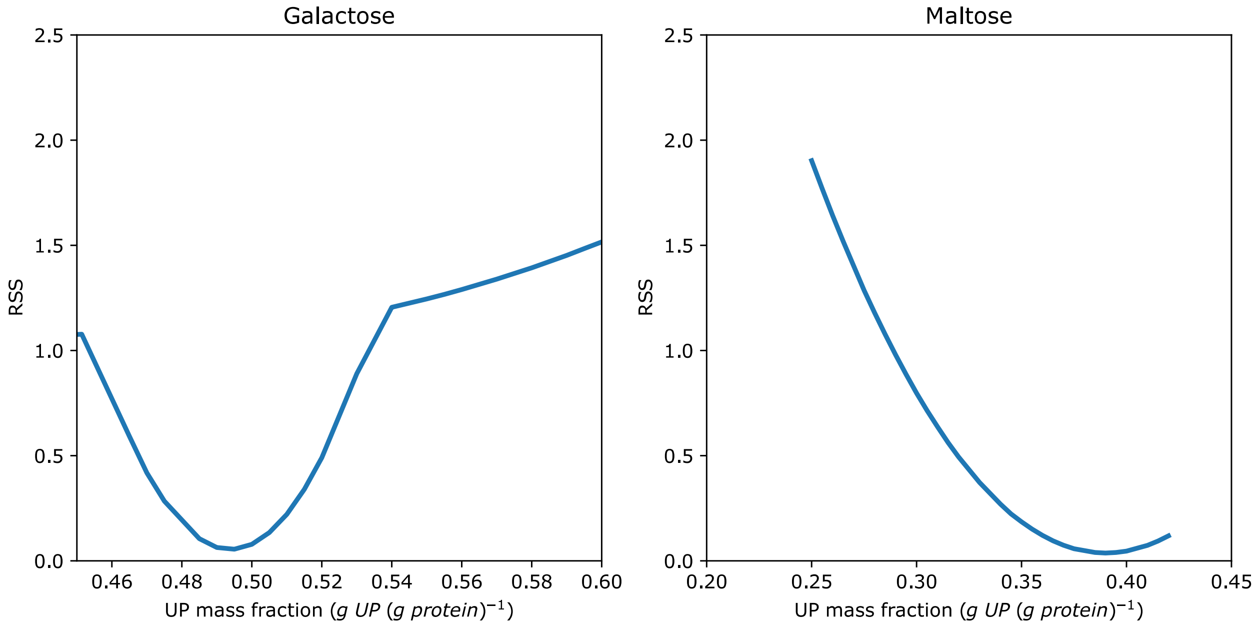

**FigSN3.** Estimation of the minimal UP mass fraction for galactose- or maltose-excess conditions.

For maltose, the goodness-of-fit measure reached values very close to zero, suggesting the flux profile well resembling that of experimental measurements. However, the fit was worse for the galactose case, even with a presence of a clear minimum of the goodness-of-fit measure in the sampled range of minimal UP values. In an attempt to better match the experimental data for galactose, we compared the fluxes of carbon source and oxygen uptake at galactose-excess conditions to glucose-limited chemostat at $D = 0.20\ h^{-1}$. The fact that consumption of the carbon source, as well as oxygen uptake on galactose was higher compared to a glucose-limited chemostat setting with comparable growth rate, suggested an additional energetic cost, which is associated with metabolizing galactose.

To simulate such an effect, we added an extra ATP maintenance reaction to the model, a “carbon-related NGAM” (cNGAM) and performed a parameter scan, similar to a previously described one. For the sake of completeness, we did an analogous scan for maltose as well. We picked cNGAM values from 0 to 5 $mmol/gDW/h$ for galactose and from 0 to 2.5 $mmol/gDW/h$ for maltose, and plotted some of the scan results in FigSN4.

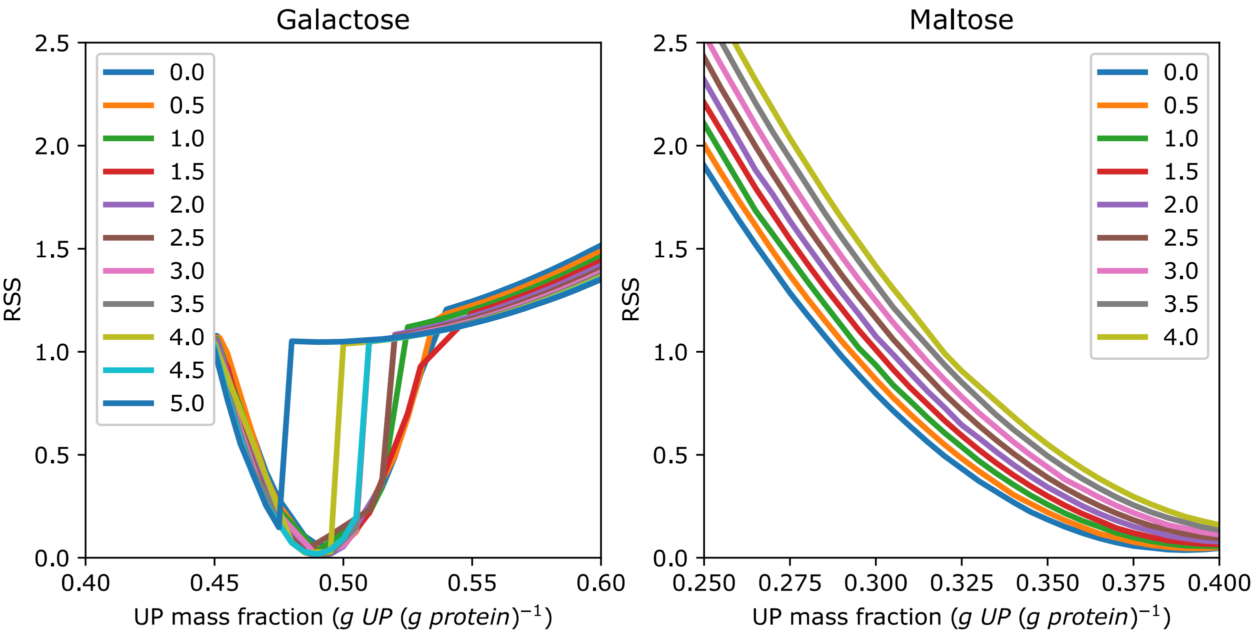

**FigSN4.** Goodness-of-fit for simulations with varying cNGAM value for galactose- and maltose-excess conditions at different UP minima.

For galactose simulations with additional energetic costs added (FigSN4, left panel), we observed a further decrease in the RSS values, suggesting an even better fit to the experimental data. To make sure this was indeed the case, we compared the predicted fluxes for galactose with the experimental data to determine the best combination of the two parameters (FigSN5).

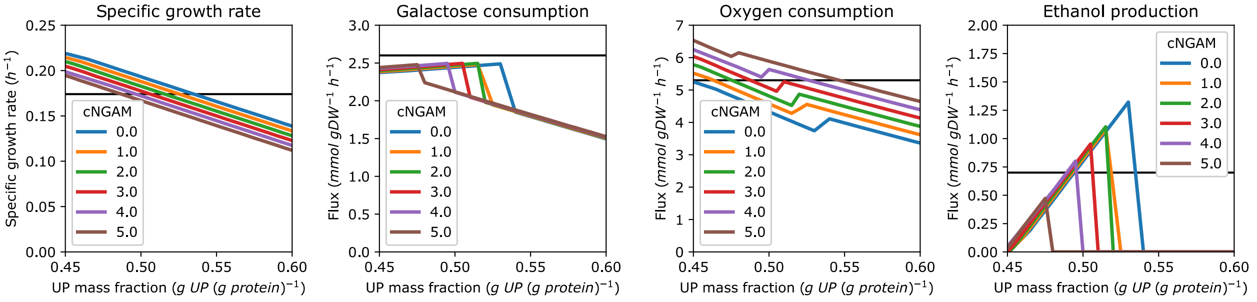

**Fig. SN5**. The flux profiles of growth on galactose with varying $cNGAM$. Lines, simulated values, horizontal black lines, experimentally measured values.

Thus, for galactose, the set of parameters, which resulted in the closest agreement to the experimental measurements (i.e. lowest RSS value), was the following: 0.49 $g\ UP/g\ protein$ and 3.0 $mmol/gDW/h$ of cNGAM. We listed the predicted specific growth rates and the exchange fluxes for galactose in SN Table 6.

For growth on maltose, additional simulations with non-zero cNGAM still suggested that the increase of the minimal UP value alone was enough to explain the experimental data. Thus we considered changing only the minimal UP value, and introduce no additional energetic cost (i.e. no cNGAM) (FigSM3, right panel). As we introduced additional proteome costs (by adjusting turnover values for maltase) alongside changing the minimal UP value, we wanted to pick a value of minimal UP, which was closest to that of the glucose-calibrated model (corresponding to 0.245 $g\ UP/g\ protein$). By setting a threshold of the $RSS < 0.25$, the first value satisfying this threshold was 0.34 $g\ UP/g\ protein$, which we accepted as sufficient (SN Table 6).

We also attempted to probe the growth on maltose with different combinations of three factors (by, e.g., implementing a change in UP minimum, non-zero NGAM but not fitted turnover value for maltase). Yet, the best agreement to the experimental data was seen in the cases where there was no additional energetic cost, some increase in the minimal UP value and the turnover value was fitted.

Thus, by using a combination of the three factors mentioned above, we show that we can describe the patterns of the metabolic fluxes for hexose sugars other than glucose by changing model parameters. We compared the predicted fractions that proteins with metabolic or unspecified function occupy with the experimental measurements, and the fractions were comparable (SN Table 7). For growth on both galactose and maltose, these changes result in two compartment-specific proteome constraints being active: maximal hexose transporter expression constraint and minimal amount of unspecified protein.

**SN Table 6.** Comparison of the experimental measurements to the predicted fluxes in the final models. Fluxes are in $mmol/gDW/h$ except for the flux through the biomass reaction (the specific growth rate, in $h^{-1}$).

| Substrate | Flux | Experimental data | Prediction after modifications | Substrate | Flux | Experimental data | Prediction after modifications |
|---|---|---|---|---|---|---|---|
| **Maltose** | $\mu\ (h^{-1})$ | 0.283 | 0.280 | **Galactose** | $\mu\ (h^{-1})$ | 0.174 | 0.182 |
| | $qS_x$ | 5.20 | 4.88 | | $qS_x$ | 2.6 | 2.47 |
| | $q_{O_2}$ | 2.70 | 4.06 | | $q_{O_2}$ | 5.3 | 5.26 |
| | $q_{EtOH}$ | 14.1 | 14.26 | | $q_{EtOH}$ | 0.7 | 0.69 |
| | $q_{CO_2}$ | 17.0 | 18.79 | | $q_{CO_2}$ | 6.0 | 6.25 |

**SN Table 7.** Comparison of proteome fractions, attributed to metabolic and unspecified proteins.

| Carbon source | Fraction of metabolic proteins + UP ($g/g\ protein$) | |
|---|---|---|
| | Experimental | Predicted with estimated UP |
| **Glucose** | 0.6809 | 0.6693 |
| **Galactose** | 0.7031 | 0.7893 |
| **Maltose** | 0.6802 | 0.7362 |

### 6. Evaluating idle ribosome content in steady-state

In this section, we follow the findings of study of (Metzl-Raz et al., 2017) to define a growth-rate dependent fraction of inactive ribosomes in pcYeast. (Metzl-Raz et al., 2017) used quantitative proteomics to determine the relationship between the specific growth rate $\mu$ and ribosomal proteome fraction $\phi_R$ for *S. cerevisiae*: $\phi_R = 0.35 \times \mu + 0.08$. This suggests that a constant fraction of ribosomes (0.08 g ribosomal proteins per gram total protein) is present at any non-zero growth rate. Our own data also suggests a presence of certain offset of ribosomal proteome fraction (main text, Fig. 1C).

Considering any growth rate, we can decompose the ribosomal fraction into the fraction of active (translating) ribosomes $\phi_R^A$ and inactive ones $\phi_R^0$: $\phi_R = \phi_R^A + \phi_R^0$. Since proteins are translated by active ribosomes only, we can consider the growth rate to be linearly dependent to the ratio of active ribosomes: $\mu = \gamma \times \phi_R^A$ (Kafri et al., 2016), where $\gamma^{-1}$ is the time required for one ribosome to replicate itself. Substituting the proteome fraction of active ribosomes into the equation for total ribosomal proteome: $\phi_R = \gamma^{-1} \times \mu + \phi_R^0$.

We can estimate $\gamma$ by considering the maximal peptide elongation rate $k_{elongation} = 10.5 \frac{aa}{s}$ (Waldron et al., 1977) and the total amino acid amount $L$ per ribosome, including assembly factors, necessary for its formation, which are 12369 and 7383, respectively (information used for calculations from (Panse & Johnson, 2010; Spahn et al., 2001)). Thus, plugging these values into the relationships, we yield $\gamma = \frac{10.5 \frac{aa}{s} \times 3600 \frac{s}{h}}{19752\ aa} = 1.9137\ h^{-1}$ and $\gamma^{-1} = 0.5225\ h$.

As we consider a steady-state molecular network, the proteome should also include the ribosome amount which *produces* ribosomes. Thus, we decompose $\phi_R^A = \phi_R^{AP} + \phi_R^{AR} \times \phi_R^0$ where $\phi_R^{AP} = \gamma^{-1} \times \mu$ is the proteome fraction of ribosomes, which produce growth-associated proteins, and another fraction $\phi_R^{AR} = \gamma^{-1} \times \phi_R^0$ is allocated towards production of ribosomes. Under zero-growth conditions, $\phi_R^{AP} = 0$, and the total ribosomal fraction reduces to: $\phi_R = \gamma^{-1} \times \phi_R^0 + \phi_R^0 = 0.08$. Solving for $\phi_R^0$ yields the steady-state zero-growth proteome fraction of inactive ribosomes of $\phi_R^0 \approx 0.053$. We further used this number to set the growth rate-independent fraction of non-translating ribosomes.
